# Generation of viral vectors specific to neuronal subtypes of targeted brain regions by Enhancer-Driven Gene Expression (EDGE)

**DOI:** 10.1101/606467

**Authors:** Rajeevkumar Raveendran Nair, Stefan Blankvoort, Maria Jose Lagartos, Cliff Kentros

## Abstract

Understanding brain function requires understanding neural circuits at the level of specificity at which they operate. While recent years have seen the development of a variety of remarkable molecular tools for the study of neural circuits, their utility is currently limited by the inability to deploy them in specific elements of native neural circuits, i.e. particular neuronal subtypes. One can obtain a degree of specificity with neuron-specific promoters, but native promoters are almost never sufficiently specific restricting this approach to transgenic animals. We recently showed that one can obtain transgenic mice with augmented anatomical specificity in targeted brain regions by identifying *cis-*regulatory elements (i.e. enhancers) uniquely active in those brain regions and combining them with a heterologous promoter, an approach we call EDGE (Enhancer-Driven Gene Expression). Here we extend this strategy to the generation of viral (rAAV) vectors, showing that when combined with the right minimal promoter they largely recapitulate the specificity seen in the corresponding transgenic lines in wildtype animals, even of another species. Because active enhancers can be identified in any tissue sample, this approach promises to enable the kind of circuit-specific manipulations in any species. This should not only greatly enhance our understanding of brain function, but may one day even provide novel therapeutic avenues to correct the imbalances in neural circuits underlying many disorders of the brain.

## Introduction

The mammalian brain is the most complex biological structure known, with innumerable distinct cell types differing in cytoarchitecture, electrophysiological properties, gene expression and connectivity^1, 2^. Recent years have seen the development of truly revolutionary molecular tools that allow neuroscientists to elucidate precise neural connectivity^3^ and monitor^4^ and manipulate^5–7^ neural activity. However, optimal use of these tools to examine the functional circuitry of the brain requires the ability to deliver them specifically to particular elements of neural circuits (i.e. neuronal cell types), rather than as a nonspecific bolus affecting all of the neurons in a brain area. The use of molecular genetics is the only method by which one can perform truly cell-type specific manipulations, as evidenced by a variety of studies using transgenic animals expressing transgenes from neuronal promoters (genomic regions just upstream of the transcriptional start site)^8, 9^. However, such approaches are limited by the fact that because genes are expressed in a variety of cell types in the brain, promoters are not specific to a single cell type. While estimates vary^10^, there are many more *cis-*regulatory elements (i.e. enhancers and repressors, distal genomic regions which help regulate where and when promoters transcribe RNA) than promoters, suggesting that enhancers may be more specific. This led us to take an approach to the generation of molecular genetic tools with augmented specificity that we call Enhancer-Driven Gene Expression (EDGE). It is based upon identifying the *cis-*regulatory elements uniquely active in particular brain regions and combining them with a heterologous minimal promoter. When we used this strategy to make transgenic mice, they were indeed significantly more specific than the presumed parent gene, often driving expression in particular sets of neurons in the brain region they were derived from ^11^.

However, while transgenic animals are powerful tools for the analysis of neural circuits, they do have some serious drawbacks. They are costly in both time and resources, can be subject to insertional effects^12, 13^, and are most practical in a limited number of species. Moreover, while they are often excellent models of disease, transgenic technologies are far from therapeutic applications. Recombinant adeno-associated viral vectors (rAAVs) can overcome many of the above issues. They can be made relatively quickly, generally do not insert into the genome, and can be used in a variety of species^14^ including humans and therefore have clinical potential as well^15–18^. However, efforts to generate cell-specific viral vectors have been largely unsuccessful to date^19–21^, with a few notable exceptions^22, 23^. This is due in large part to the fact that one can add relatively little genetic material to the relatively small genomes of rAAVs, putting most native promoters out of reach. However, most enhancers are much smaller than promoters, raising the intriguing possibility of targeting specific neuronal cell types in any species by adapting the EDGE strategy to viral vectors. Towards this end, we present results showing the generation of EDGE viral vectors specifically expressing in particular neurons of the entorhinal cortex (EC) in two different species of wildtype animals.

## Results

### Optimization of rAAV design for Enhancer-Driven Gene Expression

Because one can obtain some degree of apparent specificity with rAAVs by means other than transcriptional regulation (e.g. serotype, precise injections of minute quantities), we took steps to ensure that any observed specificity comes from the enhancer element used. First and foremost, we used an enhancer specific to a known subset of neurons in the entorhinal cortex^11^ in transgenic animals and analyzed whether the rAAV construct could match this specificity. Figure 1A shows the expression pattern obtained from crossing one of the MEC13-53 tTA driver lines to a payload line expressing the helper transgenes for the ΔG-rabies monosynaptic tracing system^11^. Expression in this cross was limited to reelin-positive (RE+), calbindin-negative (CB-) excitatory projection neurons in LII of the medial and lateral EC (i.e. stellate and fan cells, respectively^24–26^). Second, because the distinct tropisms of the various serotypes of AAV^14, 27^ is a potential confound, we used a single serotype with a wide tropism for neurons (AAV2/1, which has a mosaic capsid of serotypes 1 and 2)^28^ for all viral constructs. To ensure that the expression pattern was not due to the specifics of the injection, we injected multiple animals with the same large volume for each virus, and always used GFP-expressing rAAVs of similar titer (see Table S1) and the same coordinates in every case (see Methods for details). Figure 1B shows the result of injecting 400 nl of a virus expressing GFP from a ubiquitous CMV promoter (CMV-rAAV) into the MEC, note the widespread strong expression throughout the various layers of the entorhinal cortex, as well as subiculum and parasubiculum.

**Figure 1.**
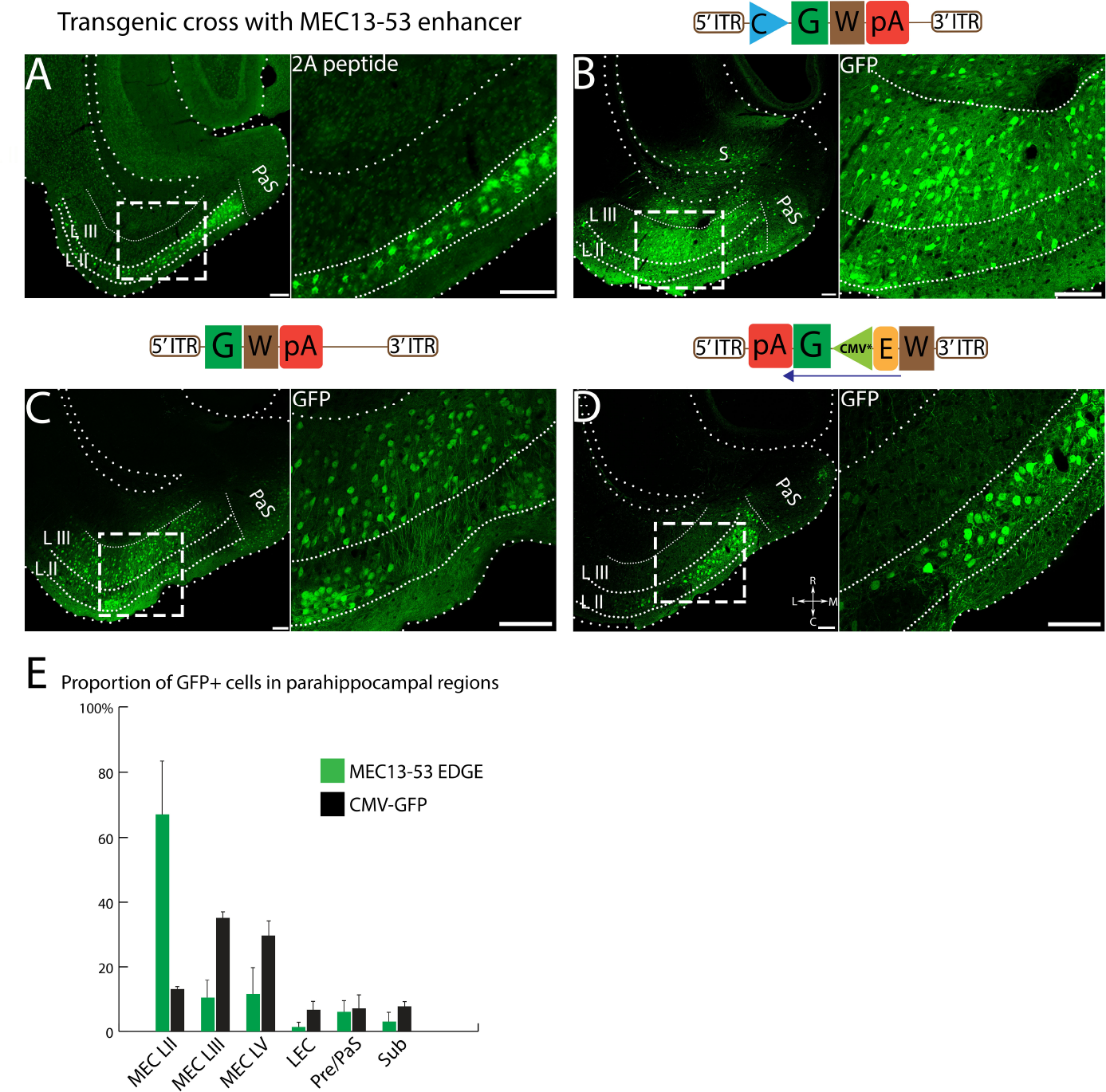
Optimization of rAAV constructs for Enhancer-Dependent Gene Expression. (A)Transgene expression in a MEC13-53 tTA × tetO-TVAG transgenic cross visualized by anti-2A immunostaining is restricted to RE+ LII projection neurons in EC^10^. (B) Injection of a nonspecific (CMV-rAAV) virus into the EC shows broad label throughout the entire region, including all layers of EC, as well as subiculum (S) and parasubiculum (PaS); (C) The same construct without a minimal promoter shows nonspecific expression throughout the region. (D) Changing the orientation of the expression cassette relative to the viral ITRs leads to a marked reduction in nonspecific expression of MEC13-53 rAAV (see inset in C and D, note that the LIII label in D is not cellular, unlike C). Images were differentially modified to best visualize the GFP expression pattern in each panel, comparisons of these images with the same post-acquisition settings are shown in figure S1. See Methods for details. (E) Proportion of GFP expressing cells in different parahippocampal regions for MEC13-53 and nonspecific CMV-rAAV. 12688 and 5063 GFP+ cells were counted from 3 mice, injected with CMV-rAAV and MEC13-53 rAAV respectively, data represented as mean ± SEM. Note that all label above background auto-fluorescence was treated as positive, even though there were two markedly distinct intensities of label (See Figure S1). Schematics of the viral designs are depicted on the top of the corresponding image. Inverted terminal repeat (ITR), woodchuck hepatitis virus post-transcriptional regulatory element (W), human growth hormone polyadenylation signal (pA), enhancer (E), Green fluorescent protein (G), cytomegalovirus promoter (C), mutated minimal cytomegalovirus promoter (CMV*). Scale bar = 100 µm.

In order to obtain viruses capable of driving expression as specific as the EDGE transgenic animals in wildtype brains one must first find a minimal viral promoter which is capable of robust expression only when paired with a heterologous enhancer. This is complicated by the fact that the viral ITRs themselves have transcriptional activity^29–31^, as can be seen by the weak nonspecific expression obtained from a GFP construct with neither a promoter nor an enhancer (Figure 1C). Note that the expression levels in 1C are far below those seen with other viruses: each panel in Figure 1 has been differentially processed to aid visualization, see Figure S1 for details. To minimize this issue, we reversed the orientation of the expression cassette relative to the ITRs such that the sense strand was under the influence of the 3’ ITR, which we attenuated by putting WPRE^32^ between the 3’ITR and the enhancer (see schematics in 1C, D). This revised design substantially reduced the background expression in other layers, enabling us to recapitulate MEC LII-specific expression in a wildtype mouse (Fig 1D) with a mutated minimal CMV promoter (CMV*)^33^. Roughly similar results were obtained with three different minimal promoters (Figure S2) but we selected CMV* for all subsequent experiments (and hereafter simply refer to the enhancer) as it was the smallest minimal promoter that yielded layer-specific EDGE with low background expression. The specificity of the expression of this virus as compared to a nonspecific CMV-rAAV virus is quantified in Figure 1E. While still clearly much more specific than the CMV-rAAV, the quantification of MEC13-53 rAAV does not look as specific as the figure panels because our conservative manual quantification for this initial characterization (see Methods) did not distinguish between weak “background” label (such as that seen without a promoter) and strong specific labelling.

### MEC13-53 EDGE rAAVs express specifically in layer II stellate cells in wildtype mice

The precise anatomical boundaries of the various layers of EC can be easily visualized by using the neuron-specific stain NeuN^34^, confirming the robust layer-specific expression of the MEC13-53 rAAV (Fig 2A). All the GFP+ cells were also NeuN positive, confirming the specificity of the virus to neurons (data not shown). Much less intense background GFP expression was observed in other layers as well in both the MEC13-53 rAAV (Figure 2A, inset) and in the rAAV backbone with the same design and minimal promoter but lacking the enhancer (Fig 2B, inset), which did not strongly label any cells. As for cell specificity, within LII of MEC there are two major classes of excitatory principal neurons, RE+ stellate cells and CB+ pyramidal cells^26, 35^, with RE label providing a sharp boundary between MEC and parasubiculum^25, 26^ (see arrows in Figure 2C, inset). We therefore performed immunohistochemical analysis comparing these markers to viral GFP and found that for the MEC13-53 rAAV, 100% (594/594) of GFP+ neurons in layer II were RE+ (Figure 2C, E), while less than 1% (5/655) of them were CB+ (Figure 2D, E). In contrast, with the ubiquitous CMV-rAAV, only 34% (319/929) of GFP+ LII neurons were RE+ while 10.5% (142/1353) were CB+. Thus, the MEC13-53 rAAV drives transgene expression specifically in a particular subset of excitatory neurons in EC of wildtype mice, i.e. RE+ EC LII neurons (stellate cells in MEC), avoiding the adjacent CB+ pyramidal cells, just as in the transgenic lines based upon the same enhancer.

**Figure 2.**
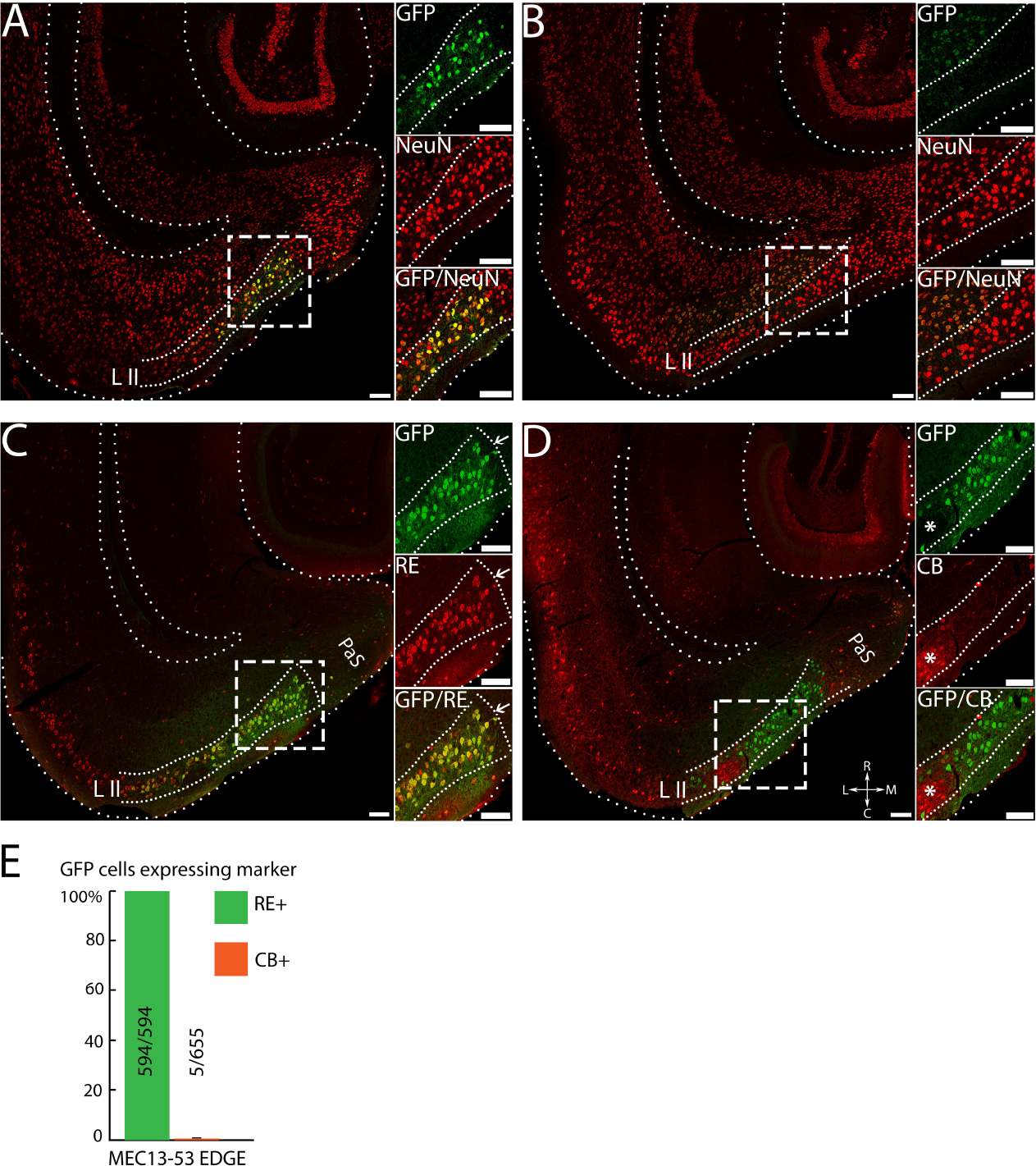
MEC13-53 EDGE rAAVs recapitulates the cell-type specificity seen in the MEC13-53 EDGE transgenic crosses in WT mice. Equal amounts of MEC13-53 rAAV (A,C,D) or CMV*-rAAV (B, i.e. identical except without any enhancer) were injected into MEC of wildtype-mice. Insets show anti-GFP (top); marker (middle); and overlay (bottom) of box in main panel, marker is NeuN in A and B. Lower panels show sections of MEC13-53 rAAV injections counterstained with anti-RE antibody (C) and anti-CB antibody (D); with a CB+ cluster (asterisks) in the insets in (D). Note the extensive co-localization of the RE stain with the GFP, the sharp delineation of the entorhinal/parasubicular boundary by both labels (arrows, C), and the exclusion of viral label from the CB clusters (asterisks, D). (E) Quantitation of results shown in C, D, showing complete overlap of GFP with cell-marker Reelin stain (green) in LII MEC, and <1% overlap of GFP with Calbindin (red), with number of cells counted. MEC-LII GFP+ cells were counted from separate RE and CB immunostained sections from 3 mice injected with MEC13-53 rAAV, data represented as mean ± SEM. Scale bar = 100 µm.

### EDGE rAAVs drive neuronal subtype-specific expression across species

While recapitulation of the expression pattern of the corresponding transgenic mouse line nicely illustrates the specificity of EDGE rAAVs, perhaps the greatest utility of viral vectors is that they can conceivably be used in wildtype animals of any species. Because evolutionary conservation is one of the hallmarks of enhancers^36^, we posited that the murine MEC13-53 enhancer might be similarly cell-specific in the rat MEC. As seen in Figure 3A, stereotactic injections of 1000 nl of the MEC13-53 rAAV into rat MEC leads to cell-type specificity as specific as that seen in the mouse (if anything, it is more specific). Figure 3A shows the MEC13-53 rAAV counterstained with NeuN, demonstrating that GFP expression is almost exclusive to MEC LII (as quantified in Figure 3E), while the few labelled neurons seen in the virus without the enhancer have no layer II specificity (Figure 3B), much as was the case in mouse (Figure 2B). Similarly, 100% (1803/1803) of GFP+ neurons in rats injected with MEC13-53 rAAVs were RE+ (Figure 3C, F), while only 1.8% (29/1589) were CB+ (Figure 3D, F), demonstrating that in both species this virus drives expression exclusively in one (RE+) subtype of EC LII excitatory neurons but not the other (CB+), even though the two subtypes are intermingled^26^. This further demonstrates that the observed specificity results from the enhancer rather than the injection site, as does the fact that this specific expression pattern is seen throughout the dorso-ventral and medio-lateral axes of the MEC (Figure S3 C). It is interesting to note that while these two markers are largely mutually exclusive, there are reports of a very small subpopulation of RE+ neurons that are also CB+^25, 37^.

**Figure 3.**
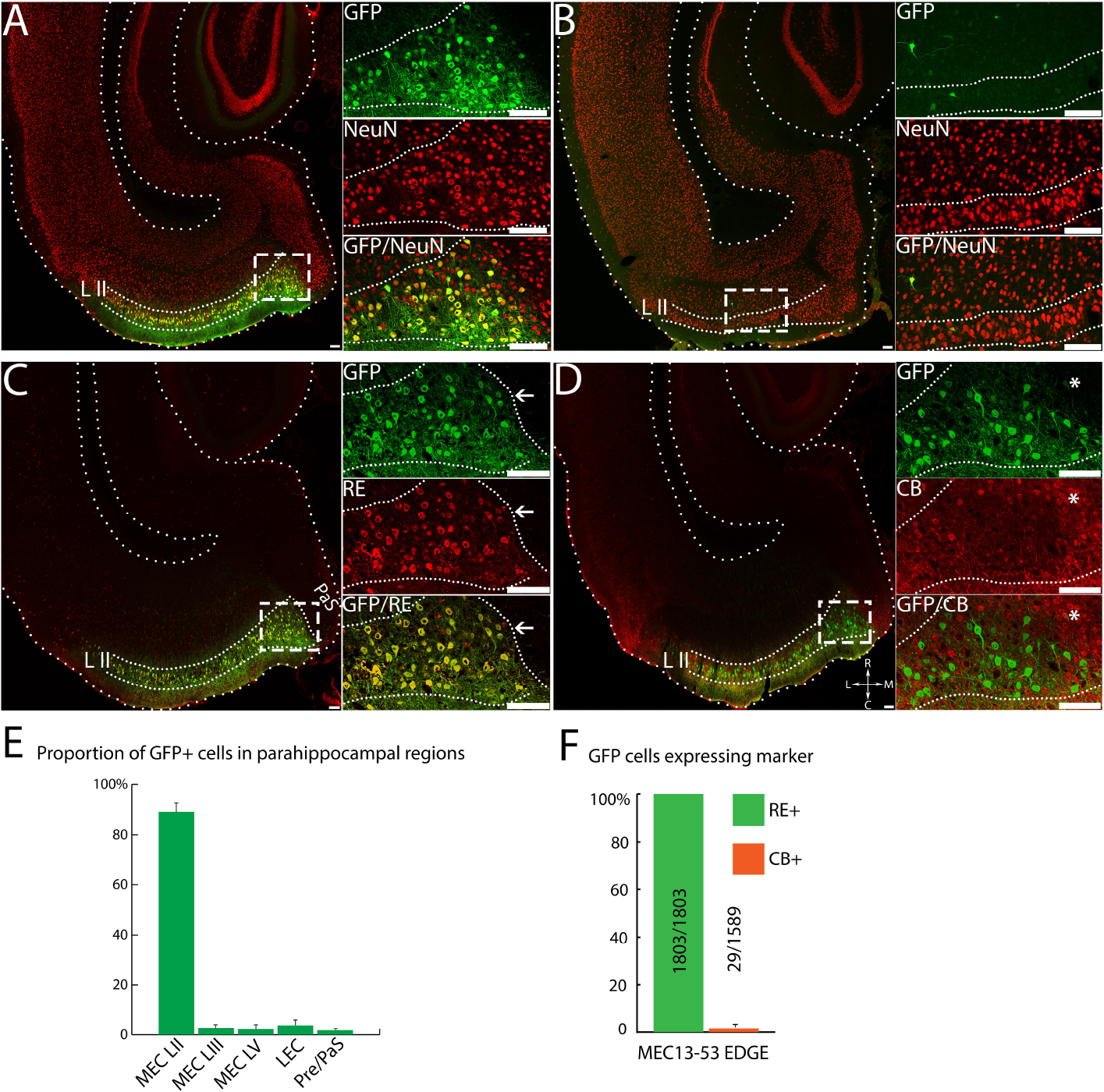
Mouse MEC13-53 rAAV is equally as specific in rat. Equal amounts of MEC13-53 rAAV (A, C, D) or CMV*-rAAV (B), i.e. identical except without any enhancer) was injected into MEC of wildtype rats. Insets show anti-GFP (top); marker (middle); and overlay (bottom) of box in main panel, marker is NeuN in A and B. Lower panels show sections of MEC13-53 rAAV injections counterstained with anti-RE antibody (C) and anti-CB antibody (D), with a CB+ cluster labelled by asterisks. Note the extensive co-localization of the RE stain with the GFP, the sharp delineation of the entorhinal/parasubicular boundary by both labels (arrows, C), and the exclusion of viral label from the CB clusters (asterisks, D). (E) Proportion of GFP expressing cells in different parahippocampal regions for MEC13-53 CMV-rAAV. (F) Quantitation of results shown in C, D, showing complete overlap of GFP with cell-marker Reelin stain (green) in LII MEC, and <2% overlap of GFP with Calbindin (red), with number of cells counted. MEC-LII GFP+ cells were counted from separate RE and CB immunostained sections from 3 rats injected with MEC13-53 rAAV, data represented as mean ± SEM. Scale bar = 100 µm.

### EDGE rAAVs recapitulate the expression pattern of their respective transgenic lines

To examine whether all EDGE rAAVs can specify gene expression in particular subsets of cells, we created EDGE rAAVs with several other enhancers. While not all enhancers that worked as transgenic lines worked in rAAVs, roughly half (Figure 4, left column) did indeed appear to recapitulate the specificity of the corresponding EDGE lines (Figure 4, right column). Remarkably, the MEC13-104 rAAV (Fig 4A) recapitulates even the sparse labeling of a small subset of LIII neurons (arrows) seen in MEC13-104, a mainly LII-specific enhancer line (Fig 4B), and the converse is true for LEC13-8, a mainly EC LIII-specific line (compare 4C to 4D). This suggests that the sparse label in the minor layer seen in the EDGE transgenic lines is specifically driven by the enhancer, rather than by random integration site-dependent mosaicism. Thus, the relative densities of the layer-specific label appear to be enhancer-specific, suggesting that the minority of cells which express outside of their primary layer may not be “noise”. Ongoing experiments seek to verify whether there are any functional distinctions between the cells labeled by the various enhancers, which often express in different subsets of what has been thought of as a single neuronal cell type, e.g. stellate cells.

**Figure 4.**
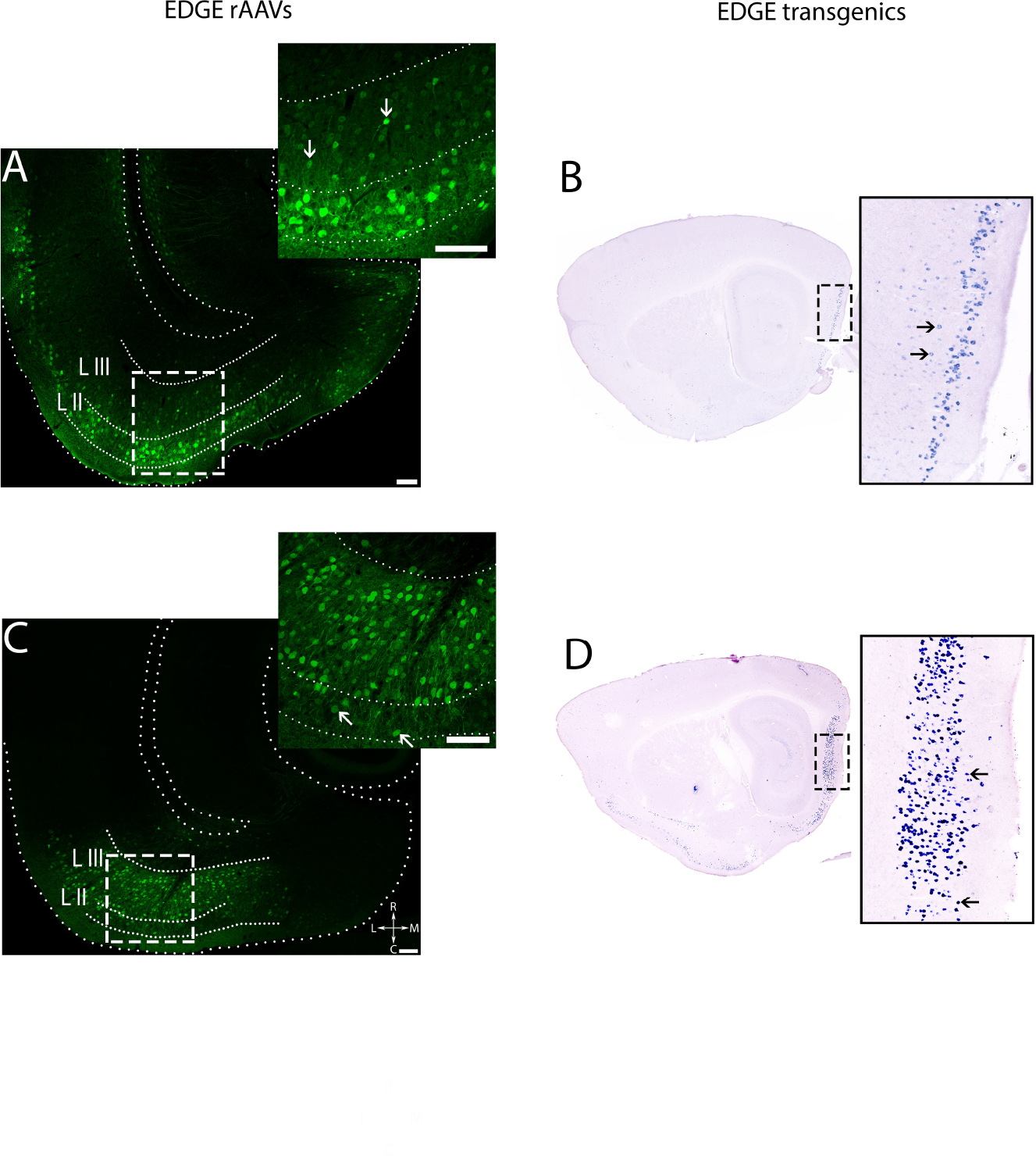
EDGE rAAVs recapitulate the distinct layer-specific expression patterns seen in EDGE transgenic mice. Comparison of expression patterns obtained by injection of EDGE rAAVs (left column) with those seen in transgenic crosses made with the same enhancers (right column). Wildtype mice were injected with 400 nl of EDGE rAAVs (A) MEC13-104 and (C) LEC13-8. Transgene expression in the corresponding EDGE transgenic crosses (B, MEC13-104 tTA × tetO-TVAG) and (D, LEC13-8 tTA × tetO-GCaMP6) visualized by ISH using the respective transgene probes. The sparse expression of the transgene in minor layers is indicated by arrows both in EDGE transgenics and viruses. Scale bar = 100 µm.

## Discussion

Our prior work showed that identification of *cis-*regulatory elements uniquely active in finely dissected cortical subregions allows one to generate genetic tools specific to cells in that subregion, an approach we call EDGE (Enhancer-Driven Gene Expression)^11^. Here we show that one can use the same approach to make rAAVs with similar specificity in both mouse and rat, provided the vector and minimal promoter’s innate transcriptional activity is minimized. This clearly cross-validates the initial identification of enhancers in our prior work^11^: while transgenic lines might show highly specific expression patterns purely due to insertional effects (though not the same patterns multiple times, as we saw), rAAVs typically do not insert into the genome^38^, so they cannot show such effects. In other words, while the precise functional significance of the enhancers presented here remains unknown, they clearly are “true” enhancers, reflecting some genetic subgroup of excitatory neurons in the entorhinal cortex of wildtype mice and rats. Taken together, these data lead to two very interesting conclusions: 1) the number of genetically-defined subgroups of neurons may be far greater than generally assumed (though this assumption has recently been challenged^39, 40^), and 2) this approach conceivably provides a path towards neuronal subtype-specific transgene expression in any species.

That enhancers drive expression similarly in both transgenic lines and viruses is not a particularly surprising result. It has been known for decades that enhancers drive cell-specific expression^41–43^. For instance, enhancers related to the 6 homeobox genes related to the fly *distal-less* gene^44^ (*Dll* in fly, Dlx in vertebrates) have been shown to play a crucial role in morphogenesis in a variety of species^45^. Due to their central role in development such homeobox genes have been under intense scrutiny by geneticists working in a variety of systems ^46–49^ for decades, leading to a highly detailed understanding of these loci and the *cis-* regulatory elements controlling their transcription. One such enhancer in the Dlx 5/6 gene cluster has been shown to be critical to the development of interneurons in particular^50^, though. A recent paper^22^ used this enhancer element in a viral vector to obtain interneuron-specific expression in a variety of species, nicely showing that enhancers can be used to drive expression in viral vectors. However, as is true for most genetically-defined enhancers active early in development, Dlx5/6 drives expression across broad classes of neurons (e.g. interneurons in general) throughout the brain, rather than to particular interneuronal subclasses and/or subregions. In addition, there are several (mostly unpublished) reports of using *cis-*regulatory elements identified by various means to drive expression in neuronal subtypes^23, 43, 51, 52^.

Thus, the most important aspect of these data is not that enhancers can work in viral vectors, it is the illustration of the promise of applying modern genomic techniques to the study of the precise neural circuitry of the vertebrate brain. The striking diversity of enhancers found in these tiny subregions of cortex (the numbers of unique enhancers in each was comparable to those found for entire organs) may indicate a similar diversity of neuronal cell types in the brain. However, the relationship between enhancers and cell types remains unclear. Indeed, the expression patterns we obtain are arguably more specific than our current understanding of neuronal cell type^1, 2^. For instance, stellate cells are a generally-accepted excitatory neuronal cell type of the medial entorhinal cortex^53^. They are characterized mainly by their stellate morphology, the fact they project to the hippocampus, and their expression of RE but not CB^25, 26^. However, we show that 2 distinct enhancers reproducibly drive expression in certain percentages of stellate cells in LII of EC, even in rAAV. The question becomes whether these enhancer-driven expression patterns reflect functionally distinct subtypes of stellate cells, or random subsets of the same indivisible cell type? In the specific case of stellate cells, a recent paper used optogenetic tagging to show that stellate cells of the MEC exhibit a variety of quite distinct receptive field properties (i.e. they can be grid cells or spatial cells or border cells, etc), suggesting that there are many functional subtypes of stellate cells^35^. More generally, the relationship between differential enhancer usage and neuronal cell types is a highly non-trivial question, not least because there is not even complete agreement even as to *how* to define neuronal cell types (though there are notable exceptions)^39, 40, 54^, let alone how many there are. There are several other interesting explanations for differential enhancer usage beyond cell type, for instance it could dictate distinct states of a single cell type. In support of this, neural activity drastically changes the chromatin landscape of the brain, including which enhancers are active^55, 56^. It will likely take years of anatomical, molecular, and physiological characterization of these tools to disentangle such questions, so for our current purposes the most important consideration is that these enhancer-based molecular genetic tools remain true to type, as appears to be the case. For instance, both the 7 MEC13-53 transgenic lines^11^ as well as the MEC13-53 rAAVs completely avoid expression in CB+ pyramidal cells, the other major kind of MEC LII excitatory neurons intercalated with stellate cells ^25, 26^, shown in Figures 2 and 3, clearly differentiating between distinct subtypes of excitatory neurons within the same cortical layer in both multiple transgenic lines and rAAVs in two species.

It should be noted, however, that specificity is never absolute, especially with viral vectors. While we obtain neuronal subtype-specific results with large injections into the entorhinal cortex (Figure S3), it is likely that any cell type in other brain regions which express the transcription factor(s) appropriate for a particular enhancer would be labeled as well. Moreover, presumably many more cells are infected than show strong GFP label, and there is a baseline level of transcription from other elements in the viral construct (i.e. the minimal promoter and the ITRs). This implies that multiple infections of an EDGE rAAV in any cell could lead to discernible nonspecific transgene expression without involvement of the enhancer at all (Figure 1C, 2B). Viral expression is thus not all-or-nothing, but the difference between background and enhancer-driven expression levels can be quite marked (Figure S1). This background expression can be quite problematic with enzymes such as recombinases, or when complementing replication-competent viruses (e.g. ∆G-rabies^57^), but is likely not an issue with transgenes whose effects vary roughly linearly with their expression levels, such as chemogenetic^6^ and/or optogenetic tools^7^.

Thus, identification of the active enhancers of a mere four cortical subregions of the mouse brain has led to a variety of viral tools for circuit analysis that appear to work across species, at least in rodents. Since one can do this on any species with a reasonably well-annotated genome, one could conceivably develop tools for anatomically specific “circuit-breaking” experiments in any species. Thus, not only will circuit-specific tools greatly facilitate our understanding of normal and pathological brain function, they could in time possibly provide novel circuit-specific therapeutic avenues. For example, it has been known for decades that preclinical stages of Alzheimer’s disease (AD) are characterized by neuronal loss and accumulation of neurofibrillary tangles in the superficial layers of trans-entorhinal cortex^58^, a region roughly equivalent to rodent MEC layer II. In addition, intracellular amyloid-β is found specifically in MEC layer II reelin-positive neurons in human AD pathology and rodent disease models^59^. Given the emerging consensus that AD may progress trans-synaptically^60, 61^, it is conceivable that one could use something like a MEC13-53 rAAV to deliver therapeutic agents directly to the presumed pre-α cells. More generally, it is possible that the reason that many neurological and neuropsychiatric disorders are resistant to drug therapy is that they are imbalances in particular neural circuits, not diseases of the entire brain. A drug having tropism for multiple circuits (as most do) would then by definition produce unwanted side effects: it may do the right thing in the right circuit, but it does the wrong thing to normal circuits. Results like those presented here allow us to hope that we may one day be able to design interventions with the required specificity to match the complexity of diseases of the brain.

---

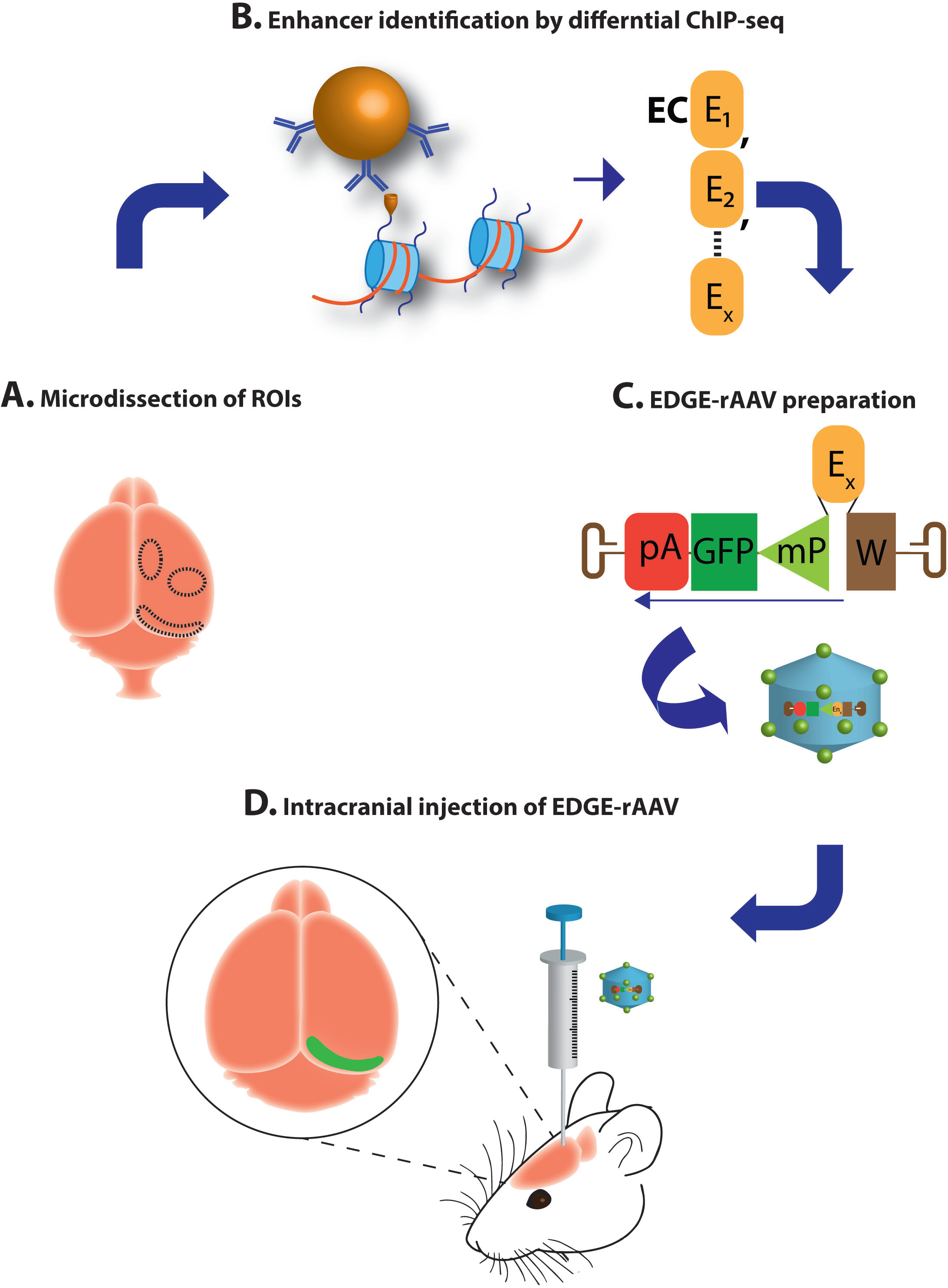
Schematic summary of region-specific transgene expression by EDGE-rAAV viruses. (A) Biological replicates of regions of interest are microdissected from C57BL/6J mouse brain. (B) H3K27ac & H3K4me2 Chip-seq and comparative analyses identify enhancers uniquely active in the region of interest. (C) Putative region-specific enhancer is cloned to rAAV vector expressing transgene under control of tight minimal promoter. (D) Purified EDGE-rAAV is stereotaxically injected into rodent brain and enhancer drives transgene expression restricted to specific brain region. Schematics of EDGE-viral design: Woodchuck hepatitis virus post-transcriptional regulatory element (W), human growth hormone polyadenylation signal (pA), enhancer (E_X_), Green fluorescent protein (GFP), minimal promoter (mP).

**Figure S1. Visualization of GFP signal depends critically on post-processing.**

The same images shown optimally in Figure 1B-D are shown with the same post-acquisition processing for the purposes of comparison of strong versus weak viral GFP expression seen with the different viral constructs. Left column is CMV rAAV, middle column is the Promoterless-rAAV, and the right column is the MEC13-53 rAAV. Top row (S1A-C) shows images at optimized settings as shown in Figure 1B-D; second row (S1D-F) shows images at optimization settings for CMV-rAAV applied to all images; third row (S1G-I) shows optimization settings for Promoterless-rAAV; fourth row (S1J-L) shows optimization settings for MEC13-53 rAAV. Note that at the settings for both CMV-rAAV and MEC13-53 the background GFP seen in the Promoterless-rAAV (SE, K) is not visible, while at the settings optimized for Promoterless-rAAV the label in the parenchyma makes it impossible to visualize individual cells in the CMV-rAAV and at low magnification makes it look like there is label in LIII in the MEC13-53 rAAV (see Figure 1D inset). Scale bar = 100 µm.

**Figure S2. Optimization of minimal promoter for EDGE-rAAV constructs**

400 nl of MEC13-53 EDGE rAAVs with the various minimal promoters (A) CMV*, (B) FGF4, (C) TK or (D) HSP68 (see Methods for details) were injected into MEC in wildtype mice. While each minimal promoter led to layer-II specificity when combined with the MEC13-53 enhancer, we chose to use CMV* because of its smaller size and limited nonspecific expression in other layers. Scale bar = 100 µm.

**Figure S3. EC LII specificity of MEC13-53 at multiple levels of caudal forebrain.**

Representative images of the GFP+ and NeuN+ neurons in horizontal sections at multiple dorso-ventral levels from, (A) a mouse brain injected with 400 nl and (C) a rat brain injected with 1000 nl of MEC13-53 rAAV. MEC13-53 drives transgene expression preferentially in MEC LII throughout dorso-ventral axis. (B) Representative images of the GFP+ and NeuN+ neurons in multiple horizontal sections in dorso-ventral axis from a mouse brain injected with CMV-rAAV. Label is throughout the layers of EC and also in subiculum. Sterotaxic coordinates were identified based on anatomical features using Paxinos G & Franklin K (for mouse brain) and Paxinos G & Watson C (for rat brain). Scale bar =100 µm.

## Methods

### Construct and rAAV preparation

All rAAV constructs were generated on backbone plasmid pAAV-CMV-MCS-WPRE-hGH PolyA (modified by cloning WPRE after the MCS in pAAV-MCS, Agilent, USA). Control rAAV constructs were generated as follows: for ubiquitously expressing rAAV construct (CMV-rAAV) without a region-specific enhancer was generated by cloning the enhanced GFP (GFP) into the MCS of the pAAV-CMV-MCS-WPRE-hGH PolyA. For the promoterless construct for testing the transcriptional activity of ITRs, CMV promoter was removed from CMV-rAAV. To synthesize EDGE rAAV constructs, the CMV promoter, MCS and hGH PolyA sequences except WPRE were removed from pAAV-CMV-MCS-WPRE hGH PolyA. An expression cassette consisting of a hybrid promoter (composed of a region specific enhancer and minimal promoter), GFP and PolyA sequence were then subcloned into the plasmid in reverse orientation relative to the ITRs, to circumvent any promoter activity from the 5’ITR. The expression cassette in the reverse orientation was cloned into the plasmid upstream of the WPRE which thus minimized the promoter activity from the 3’ITR. Various EDGE rAAV constructs with the revised design were generated by cloning murine enhancers obtained from our initial enhancer screen such as MEC-13-53, MEC-13-104 or LEC-13-8 and different minimal core promoters such as a variant of CMV (CMV*)^33^, FGF4^62^, HSV-TK or HSP68^11, 36^ into the expression cassette in the reverse orientation. Sequences of the EDGE rAAVs, the region specific enhancers and the minimal promoters used in the study are given below. Plasmids were maintained in the Stbl3 *E. coli* strain (ThermoFisher, USA) to avoid ITR-mediated recombination. Enhanced GFP, WPRE, LEC-13-8 and minimal promoters were synthesised by Genscript, USA. Positive clones were confirmed by restriction digestion analyses and subsequently by DNA sequencing. Endotoxin-free plasmid maxipreps (Qiagen) were made for rAAV preparations.

EDGE rAAVs were packaged in AAV serotype 2/1 (having a mosaic of capsid 1 and 2)^28^ using Heparin column affinity purification^63^. Specifically, a pAAV construct generated as described above with AAV helper plasmids encoding the structural elements, were transfected into the AAV-293 cell line (Agilent, USA). The day before transfection, 7 × 10^6^ AAV-293 cells were seeded into 150 mm cell culture plates in DMEM containing 10 % fetal bovine serum (ThermoFisher, USA) and penicillin/streptomycin. Co-transfection of plasmids such as pAAV-containing the transgene, pHelper, pRC (Agilent, USA) and pXR1 (NGVB, IU, USA) was carried out next day. After 7 hours, the medium was replaced with fresh 10 % FBS-containing DMEM. The AAV-293 cells were cultured for two days following transfection to allow rAAV synthesis to occur. The AAV-293 cells filled with virus particles were scraped from the cell culture plates, then isolated by centrifugation at 200 × g. The cell pellet was then subjected to lysis using 150 mM NaCl-20 mM Tris pH 8 buffer containing 10 % sodium deoxycholate. The lysate was treated with Benzonase nuclease HC (Millipore) for 45 minutes at 37°C. Benzonase-treated lysate was centrifuged at 3000 × g for 15 mins and the clear supernatant then subjected to HiTrap^®^ Heparin High Performance (GE) affinity column chromatography using a peristaltic pump^63^. The elute from the Heparin column was concentrated using Amicon Ultra centrifugal filters (Millipore). The titer of the resultant viral stock was determined by quantitative PCR as approximately 10^11^ infectious particles/ml.

### Titration of the rAAVs

The titration of the rAAVs prepared for the study was carried out by quantitative PCR^64, 65^ using Power SYBR™ Green PCR Master Mix (ThermoFisher, USA), the following primers were used for GFP; forward primer-5’-AGCAGCACGACTTCTTCAAGTCC and reverse primer 5’-TGTAGTTGTACTCCAGCTTGTGC (modified protocol from Addgene, USA). A known concentration (2 × 10^9^ molecules/µl) of a pAAV construct containing the GFP sequence was used for generating the standard curve. 5 serial dilutions of plasmid from 2 × 10^8^ to 2 × 10^5^ were made in PCR grade water for creating the standard curve. The purified rAAVs were treated with DNase I (ThermoFisher, USA) at 37°C for 30 minutes, to eliminate any contaminating plasmid DNA carried over from the rAAV production process. DNAse-treated rAAVs were serially diluted for the qPCR titration (from 1:20 to 1:2500) in PCR grade water. A mastermix of the reagents for the qPCR was prepared consisting of the SYBR Green PCR Master Mix, the primers and PCR-grade water. 5µl each from the standards and the rAAV dilutions along with 15µl of mastermix were subjected to qPCR at 95°C 10 min / 95°C 15 sec / 60°C 1 min/ repeat 40×/ melt curve using StepOne machine (Applied Biosystems, USA). Data analyses were performed by StepOne2.3 software and by Microsoft excel.

### Rodent Details

Experiments were carried out using C57BL/6J mice obtained from Jackson laboratory, USA and Long Evans rats (Charles River, USA). EDGE transgenic crosses were created at the Kavli Institute for Systems Neuroscience and Centre for Neural Computation as described^11^. All experiments were conducted in compliance with protocols approved by the Norwegian Food Safety Authorities and European Directive 2010/63/EU. All mice and rats were housed in enriched environment cages in a 12 hr light/dark cycle with food and water ad libitum.

### Stereotaxic Injections and Perfusions

For rat experiments, the rAAVs were stereotactically injected into three-four month old Long-Evans rats. Injections were performed with 1 µl rAAV at a titer of ~1 ×10^11^ infectious particles/ml, into the MEC of the rats. The rats were deeply anaesthetized with isoflurane gas (induction with 5 % isoflurane (v/v), maintenance at 1 % isoflurane (v/v), airflow of 1200 ml/min). To maintain the body temperature of the animal, a heating pad at 37°C was used.

Rats were injected subcutaneously with buprenorphine hydrochloride (Temgesic®, Indivior) and Metacam^®^ (Boehringer Ingelheim Vetmedica) at the prescribed dosage. Local anaesthetic Bupivacaine hydrochloride (Marcain^TM^, AstraZeneca) was applied at the place of incision. The head was fixed to the stereotaxic frame with ear bars, and the skin at the incision site was disinfected with 70 % ethanol and iodine before the incision was made using a sterile surgical scalpel blade. After incision, the mouthpiece and ear bars were adjusted so that bregma and lambda were aligned horizontally. Mediolateral coordinates were measured from the mid-sagittal sinus, anterior-posterior coordinates were measured from posterior transverse sinus, and dorso-ventral coordinates were measured from the surface of the brain. A craniotomy was made around the approximate coordinate, and precise measurements were made with the glass capillary/Hamilton needle (HAMI7762-06) used for virus injection. Coordinates for rat injections were 4.6 mm lateral, 0.2 mm anterior to the posterior transverse sinus and 2.6 mm deep, with the glass capillary/needle lowered at 10° pointing towards the nose. A single injection of 1 µl virus was conducted at a speed of 100 nl/min using a nanoliter injector (Nanoliter2010, World Precision Instruments, Sarasota, FL, USA), controlled by a microsyringe pump controller (Micro4 pump, World Precision Instruments). After completion of the injection, the capillary was retracted after a 10 minute delay, to give the virus time to diffuse. Finally, the wound was rinsed with saline and the skin was sutured. The animals were left to recover in a heating chamber, before being returned to their home cage. Next day Metacam was administered orally and their health was checked daily.

For mouse experiments, 10-15 week-old adult C57BL/6J mice were anaesthetized with isoflurane (induction with 5 % isoflurane (v/v), maintenance with 1 % isoflurane (v/v), airflow of 1200 ml/min). After applying the local analgesic Marcain (40 µl, 0.25 mg/ml, SC), the global analgesic Temgesic (0.03 mg/ml, 100-150 µl per mouse dependent on bodyweight, SC), and Metacam (2.5 mg/ml, 100-150 µl per mouse dependent on bodyweight, SC) the head was fixed in a stereotaxic frame. Subsequently the skull was exposed by a single incision of the scalp, craniotomies were made approximately 5 mm posterior and 3.3 mm lateral of the bregma. Then, the virus solution was injected at a location 0.3-0.5 mm anterior to the transverse sinus and at a depth of 1.8-2.0 mm from the brain surface. Unless otherwise stated, all injections were bilateral injections of 400 nl rAAV injected at a rate of 50 nl/min. Mice were given a second post-operative injection of Metacam the next day, and their weight was monitored until stable.

After 4 weeks, the rodents were sacrificed. Rodents were anaesthetized with pentobarbital and perfused transcardially with freshly prepared 0.9 % saline followed by 4 % paraformaldehyde in 0.1 M Phosphate buffer (pH 7.4) with 0.9 % saline. The brains were stored in 4% PFA overnight before being transferred to 30% sucrose solution for approximately two days.

### Immunostaining

Horizontal rat brain sections of 50 µm were prepared using a sliding microtome at −30°C. Brain sections were stored at −20°C in 0.1 M phosphate buffer containing 25 % glycerin and 30 % ethylene glycol. Multiple labelling of free-floating sections was carried out as briefly described. Every sixth section in the series was selected for immunostaining and washed in phosphate-buffered saline (PBS). Sections were permeabilized and blocked for 1 hour at room temperature using PBS containing 0.1 % Triton X-100 and 3 % normal donkey serum, or, when staining for reelin and calbindin 0.5 % Triton X-100 and 5 % goat serum and when staining for NeuN 0.3 % Triton X-100 and 3 % BSA (PBS++). Sections were subsequently incubated with primary antibodies in PBS++ at 4°C for two days with mild shaking. PBS-washed sections were incubated for 2 hours at room temperature with secondary antibodies diluted in PBS++ (or PBS containing Triton X-100 without serum/BSA).

Solution containing 2.5 % 1,4-diazabicylo[2.2.2]octane/polyvinyl alcohol (DABCO/PVA) was used to mount the sections in Polysine slides (Menzel-Glaser, ThermoFisher, USA). Antibodies used were rabbit anti-GFP (ThermoFisher/Life technologies, USA A11122, 1:500), mouse anti-Reelin (Merck Millipore MAB5364, 1:1000), mouse anti-Calbindin (Swant D-28 K, CB300, 1:5000), mouse anti-NeuN (Merck Millipore MAB377, 1:1000) and rabbit anti-2A peptide (Merck Millipore ABS31, 1:2000). All corresponding secondary antibodies were from ThermoFisher/Life technologies or Jackson ImmunoResearch laboratories, USA, 1:400.

### Confocal Imaging and Analysis

Sections were imaged using a confocal microscope (Zeiss LSM 880, Zen 2012 software) with either Plan-Apochromat 40×/1.4 Oil DIC M27 oil immersion or Plan-Apochromat 20×/0.8 air immersion objectives. Imaging and analyses were conducted by the same person to control inter-analyst variation. Representative images for figures were processed equally using Zen 2012 software and were created in Adobe photoshop. Acquisition of images were done at similar capture settings in confocal microscope. For figure S1 the post-acquisition modifications were carried out similarly using Zen 2012 software. For figure 1 (B-D) images were modified differently to visualize the distinct GFP expression pattern (details in Figure S1). For the post-acquisition processing, confocal czi. images were opened in Zen 2012, and visualized with intensity range indicator. The intensity histogram for the green channel was altered until the optimum intensity was visualized (as displayed by the range indicator). Identical changes in the intensity levels were applied to the other confocal images captured using the same settings for comparison (Figure S1).

The quantification of GFP+, NeuN+, RE+ or CB+ cells was performed out manually using Zen 2012 software. Approximately ten 50 µm thick horizontal sections were selected from the dorso-ventral axis (−1.5 mm to −4 mm from Bregma) per brain. Counts were carried out on the confocal images (.czi format) of rAAV injected brain sections immunostained for GFP or respective markers (NeuN, RE, CB). For the layer specificity analyses shown in Figure 1E, 12688 and 5063 GFP+ cells were counted from approximately 30 sections from 3 mice, injected with CMV-rAAV and MEC13-53 rAAV respectively. Any fluorescent signal greater than background auto-fluorescence was considered as positive, even though there was often a baseline transcription from the rAAV promoter construct versus the enhancer-assisted (EDGE) signal, which was typically orders of magnitude greater. For analyses in rat brains (Figure 3E), 7316 GFP+ cells from sections were counted from 75 horizontal sections from 5 separate hemispherical injections of MEC13-53 rAAV in 3 rats (−2 mm to −6 mm from Bregma). For the quantification of MEC-layer II EDGE GFP cells expressing cell-markers, 594 (RE co-immunostained) and 655 (CB co-immunostained) GFP+ cells were counted from 3 separate MEC13-53 rAAV injected mice (Figure 2E). For similar analyses in rats (Figure 3F), 1803 and 1589 MEC-LII GFP+ cells were counted from separate RE and CB immunostained sections from 5 different hemispherical injections of MEC13-53 rAAV (we analysed 30 sections for RE and 28 sections for CB). Analyses were done in Microsoft excel and graphs were made in Adobe illustrator.

### In situ hybridization

Mice were perfused transcardially with 4% paraformaldehyde (PFA) in RNAse free PBS. The brain was extracted and stored in 4% PFA overnight before being transferred to 30% RNAse free sucrose solution for approximately two days. The brain was then sectioned horizontally in 30 µm thick sections and divided into a set of approximately 6 series and stored in a −80°C freezer. A series was then thawed before use. To stain transgenes TVAG or HM3, 30μm thaw-mounted sections were hybridized overnight at 62°C with a DIG-labelled riboprobe for TVAG or HM3 (approximately 1:500; Roche, Cat. 11277073910) and then incubated in Blocking solution (600µl MABT, 200µl sheep serum, 200µl 10% blocking reagent (Roche, Cat. No. 11 096 176 001)) for 2-3 hours at room temperature. Slides were drained of the blocking solution, and antibody solution (1:5000 dilution of sheep anti-dig alkaline-phosphatase (AP) in blocking solution) was added to the slides. The slides were transferred back to the Perspex box and incubated at room temperature overnight. 4g of polyvinyl alcohol (Mol. Wt. 70000 – 100000) were transferred to a 50ml polypropylene centrifuge tube, and AP staining buffer (100mM NaCl, 50mM MgCl_2_, 100mM Tris pH 9.5, 0.1% Tween-20) was added till the total volume of the solution was 40 ml. The solution was shaken to dissolve the solid material, and further heated in a water bath. When the solution was clear, it was cooled down to 37°C. The slides were washed 5 times in MABT in room temperature for 4 min each wash. Further, the slides were washed 2 times in AP staining buffer for 10 min in room temperature while the slides were shaken. NBT (Nitroblue tetrazolium chloride, Roche. Cat. No. 11 383 213 001. 140 µl/40ml), BCIP (5-Bromo-4-chloro-3-indolyl-phosphate, 4-toluidene salt, Roche, Cat. No. 11 383 221 001. 105 µl/40 ml) and Levamisole (Vector, Cat. No. SP-5000. 3.2ml/40ml) were added to the polyvinyl alcohol solution, and mixed well before transferring it to a Coplin jar together with the slides. The slides were incubated at 37°C for 5 hours. After incubation, the slides were washed 2 times in PBS + 0.1% Tween-20 to stop the staining reaction. The slides were further washed 2 times in ddH_2_O and left air dry at room temperature for overnight. The slides were cleared in xylene and coverslipped. The stained sections were imaged using automated Axio Scan. Z1 scanner (Zeiss), Zen 2.3 software with transmitted white light as the light source.

## Supplemental information

**Table S1.**
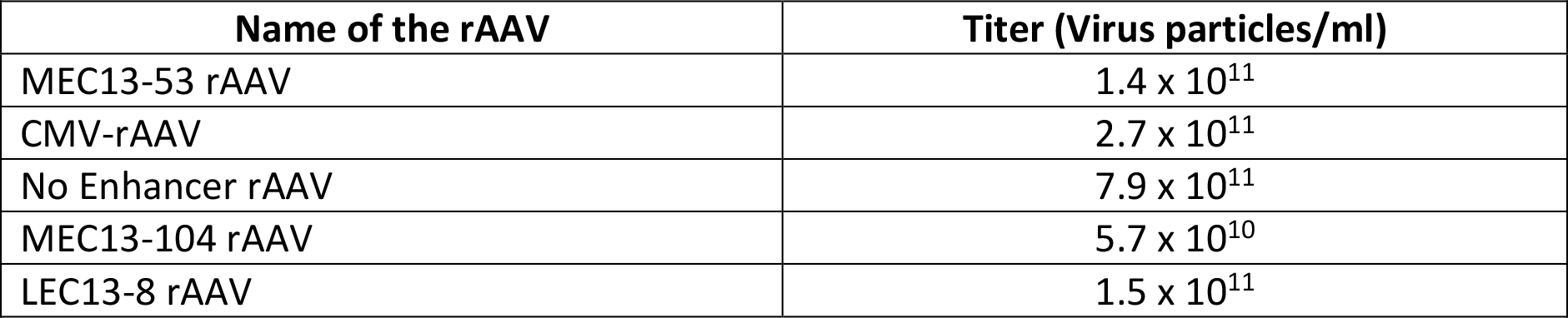
Titers of the important rAAVs synthesized for the current study. Quantitative PCR was carried out for the titration of the rAAVs. See the methods sections for detailed protocol.

| Name of the rAAV | Titer (Virus particles/ml) |
| --- | --- |
| MEC13-53 rAAV | $1.4 \times 10^{11}$ |
| CMV-rAAV | $2.7 \times 10^{11}$ |
| No Enhancer rAAV | $7.9 \times 10^{11}$ |
| MEC13-104 rAAV | $5.7 \times 10^{10}$ |
| LEC13-8 rAAV | $1.5 \times 10^{11}$ |

## Sequences of the minimal promoters used in the study

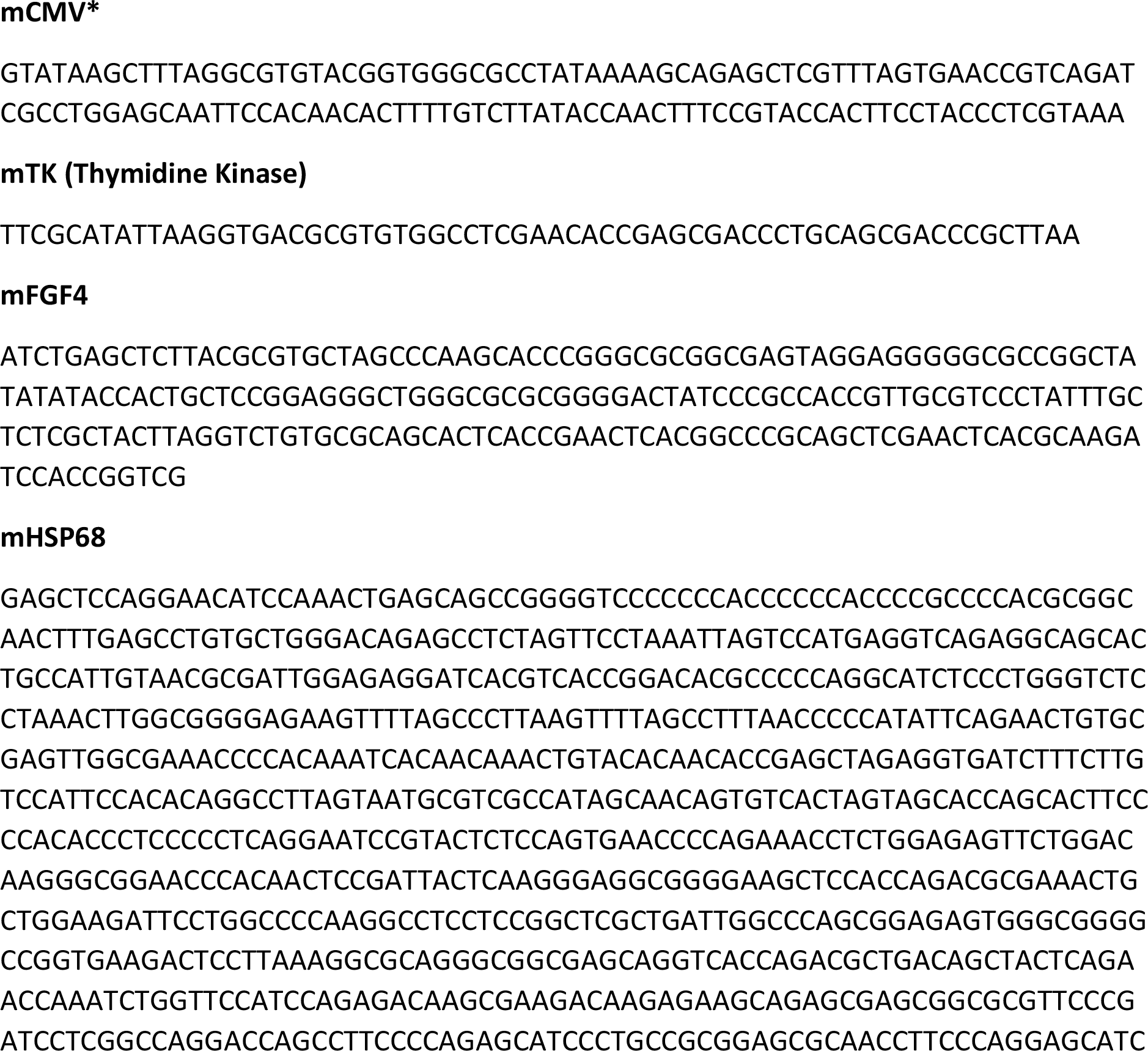
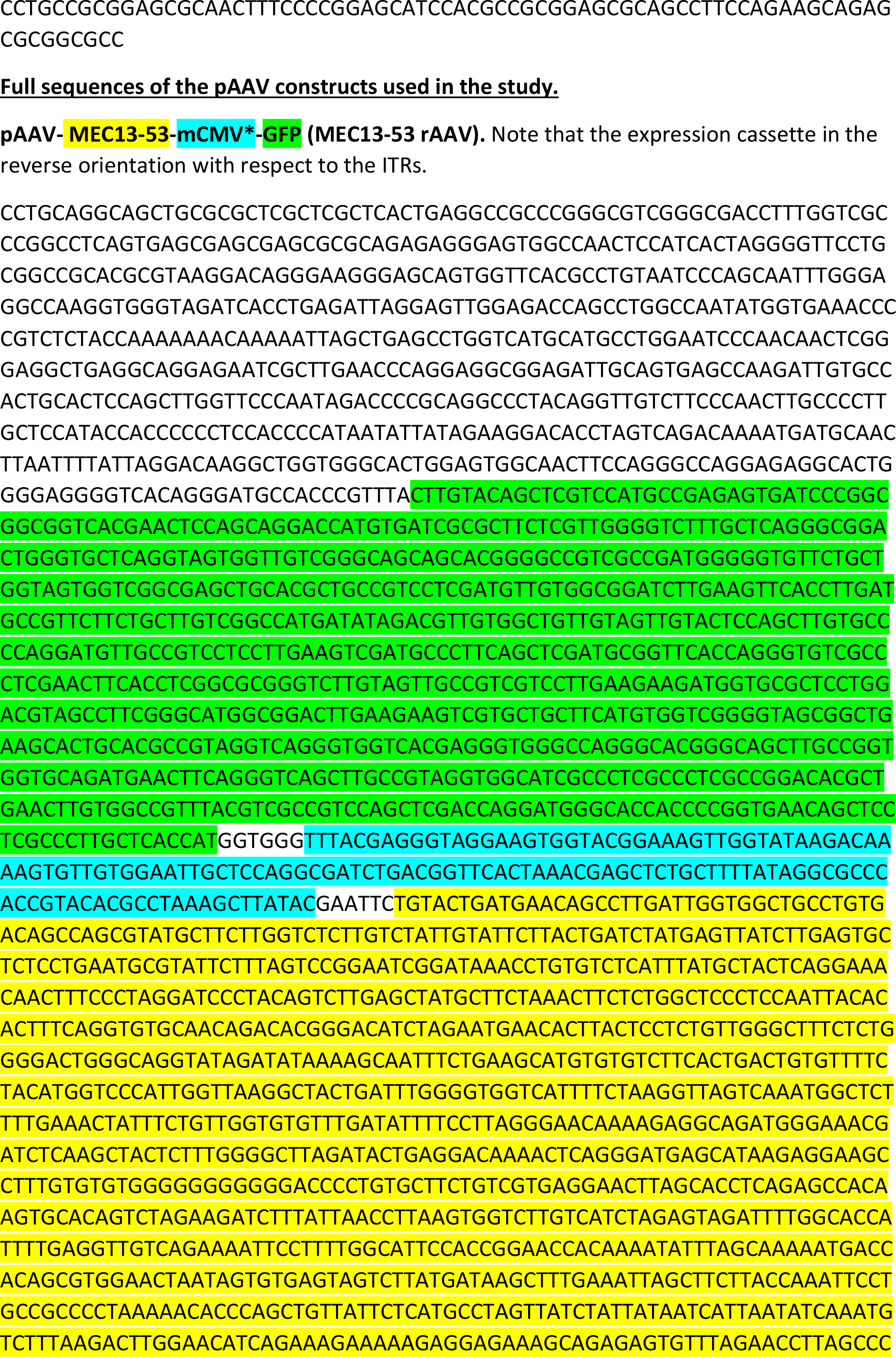
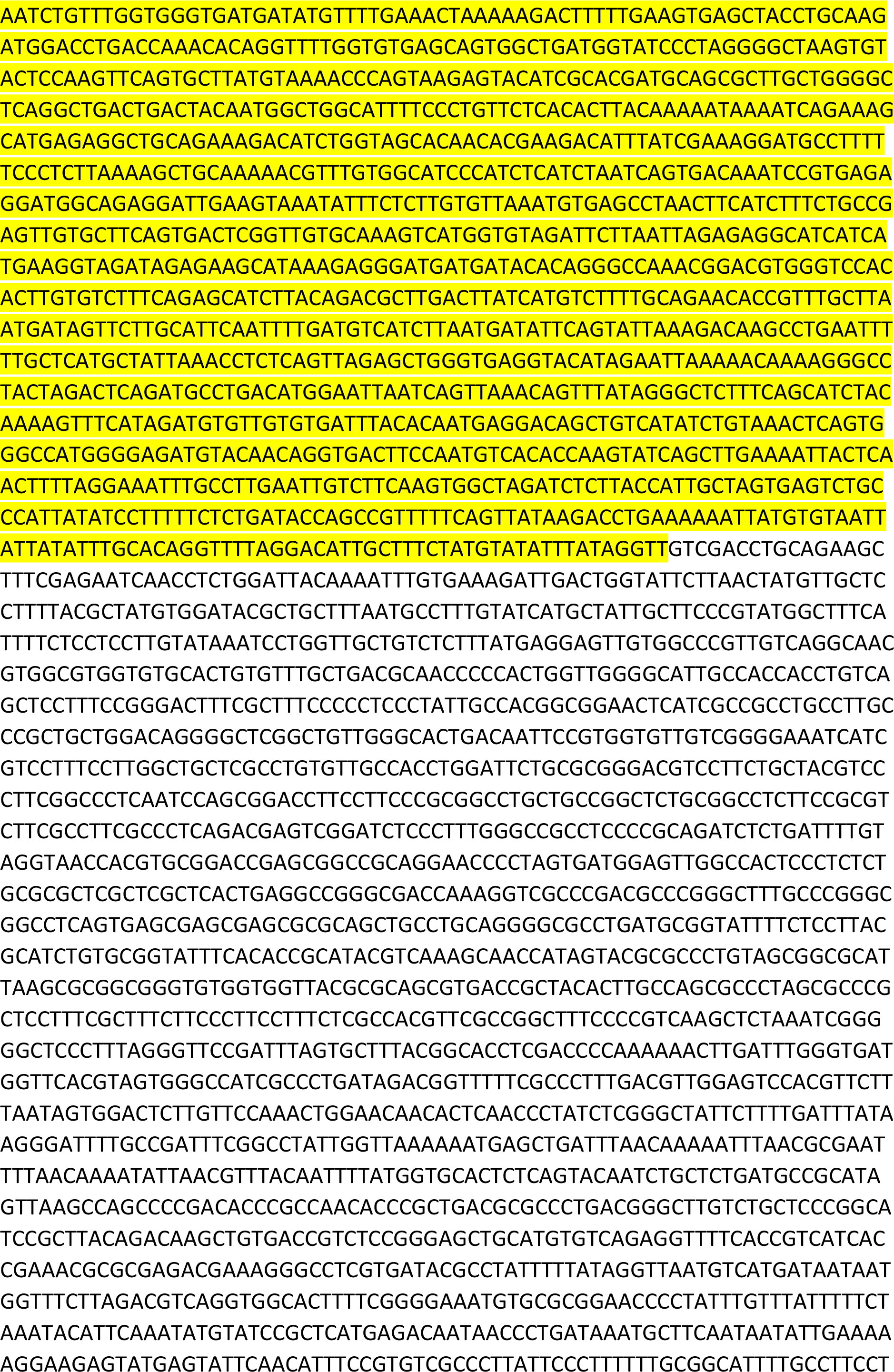
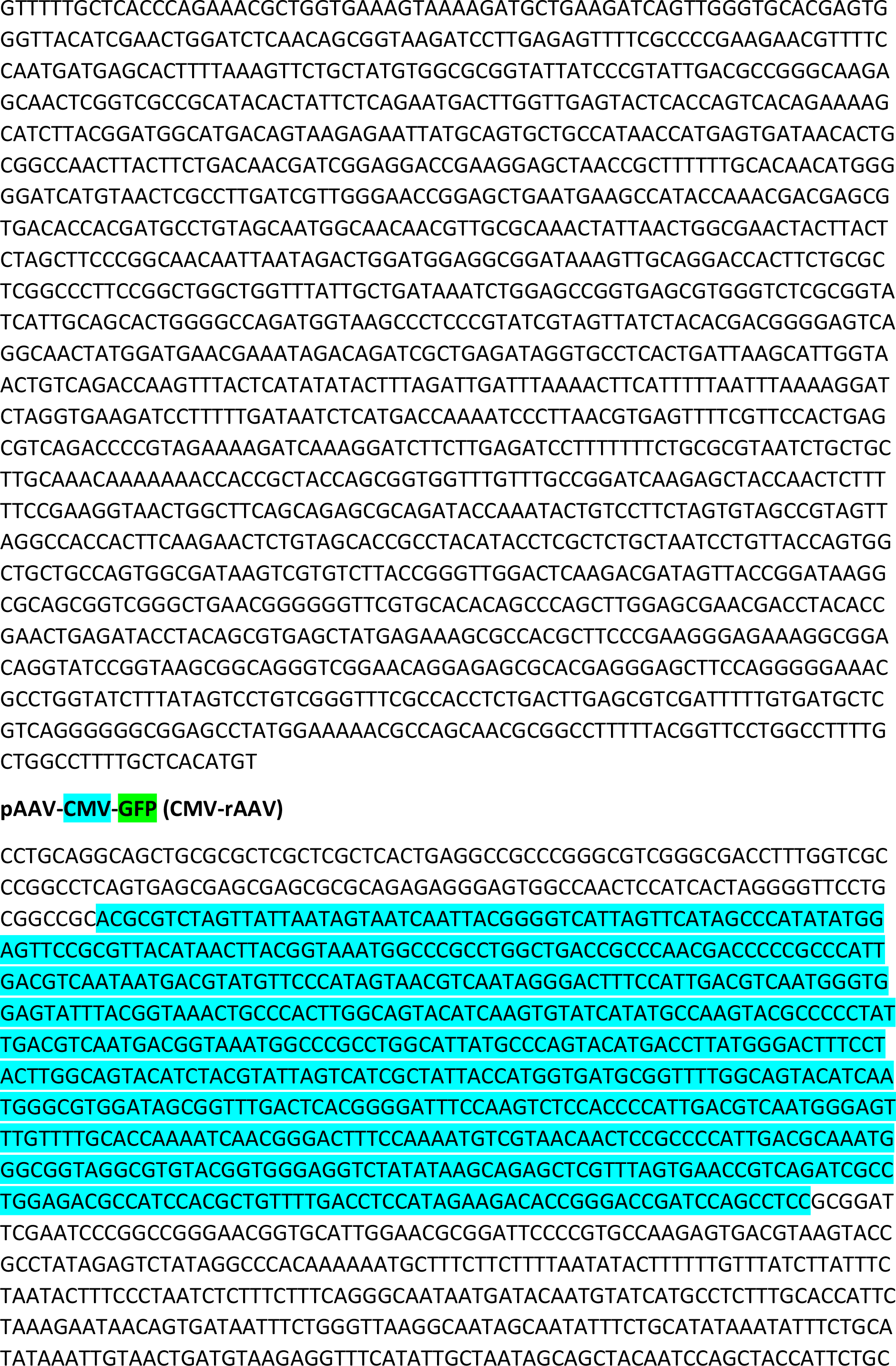
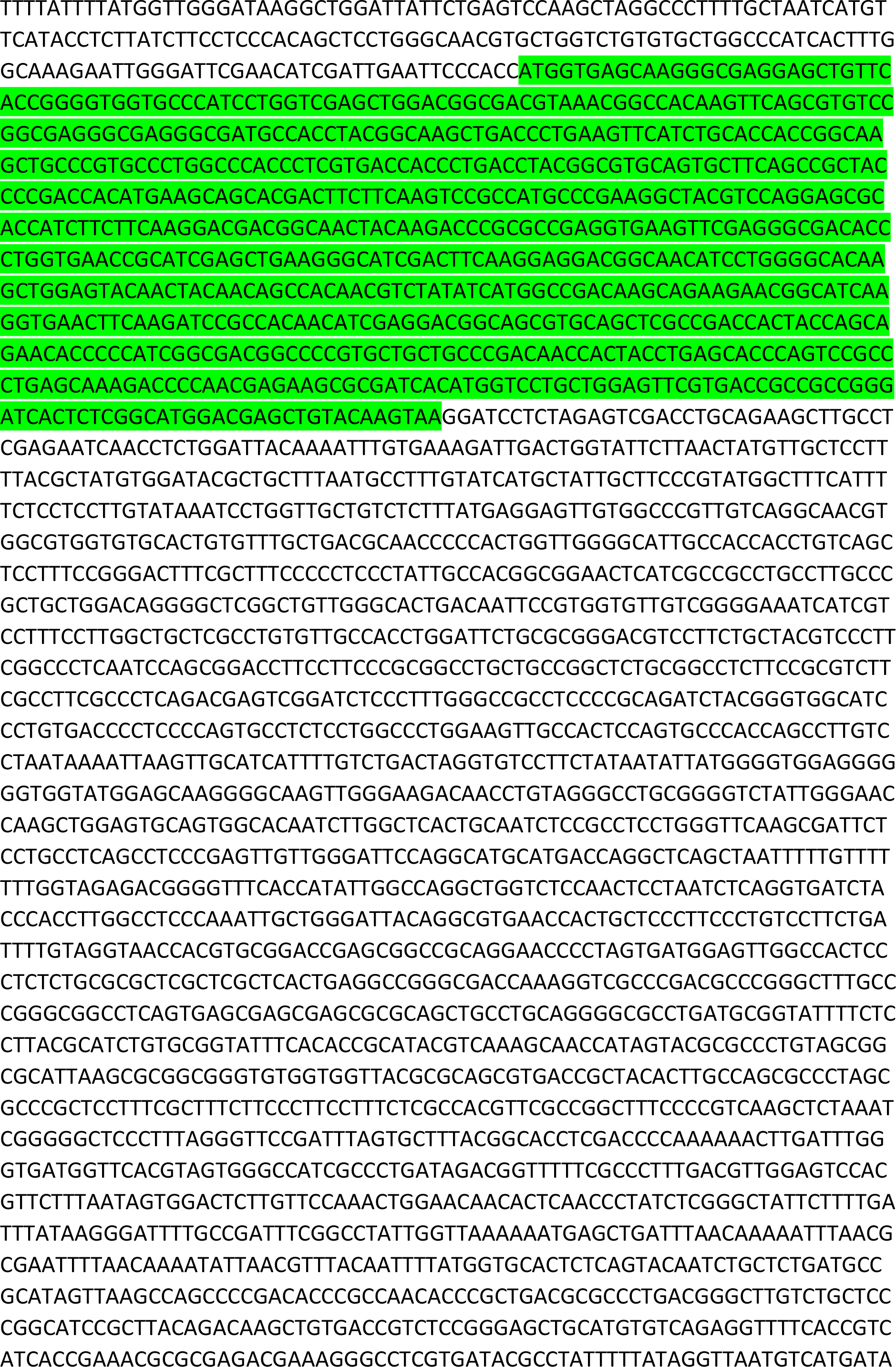
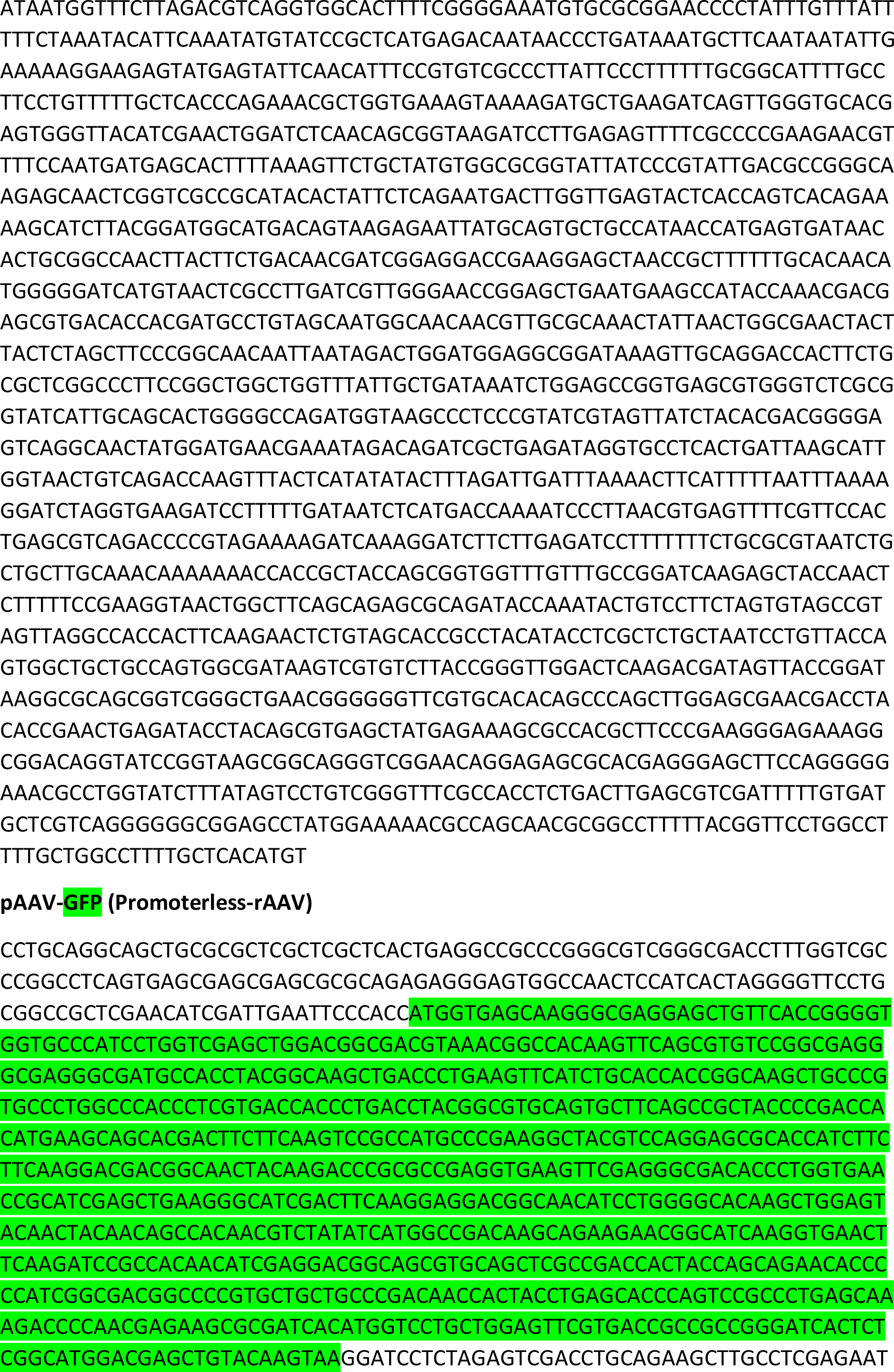
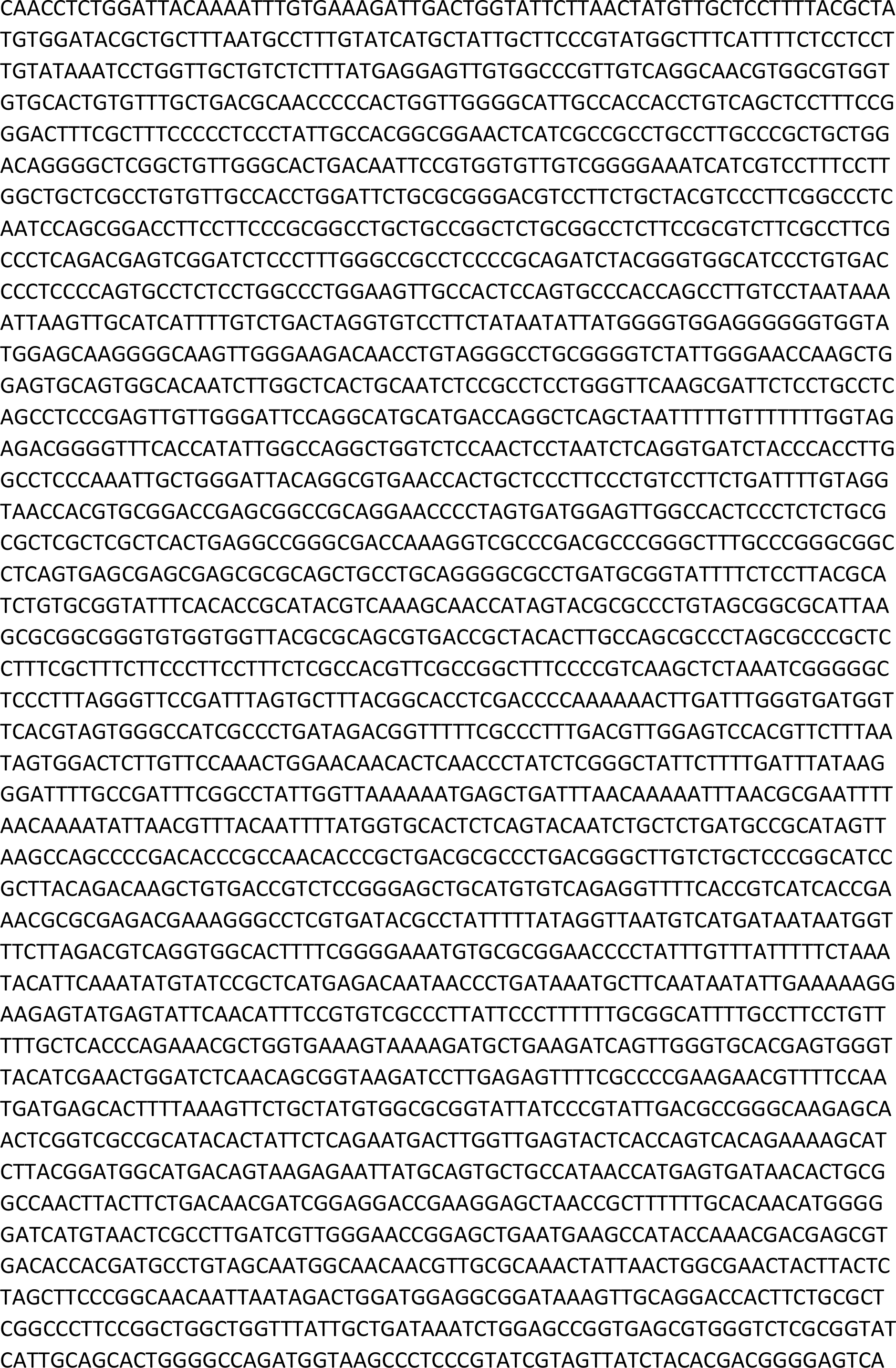
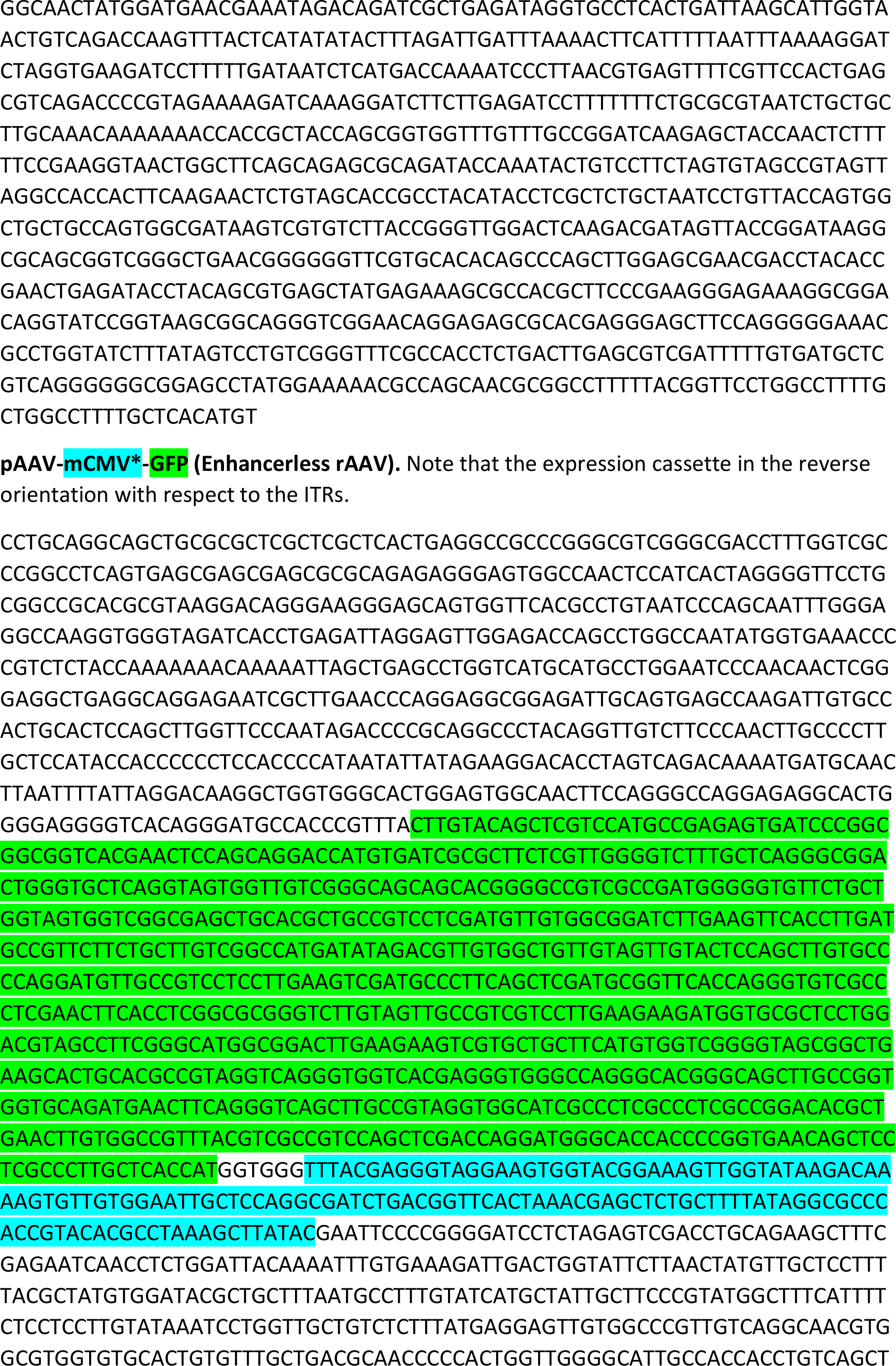
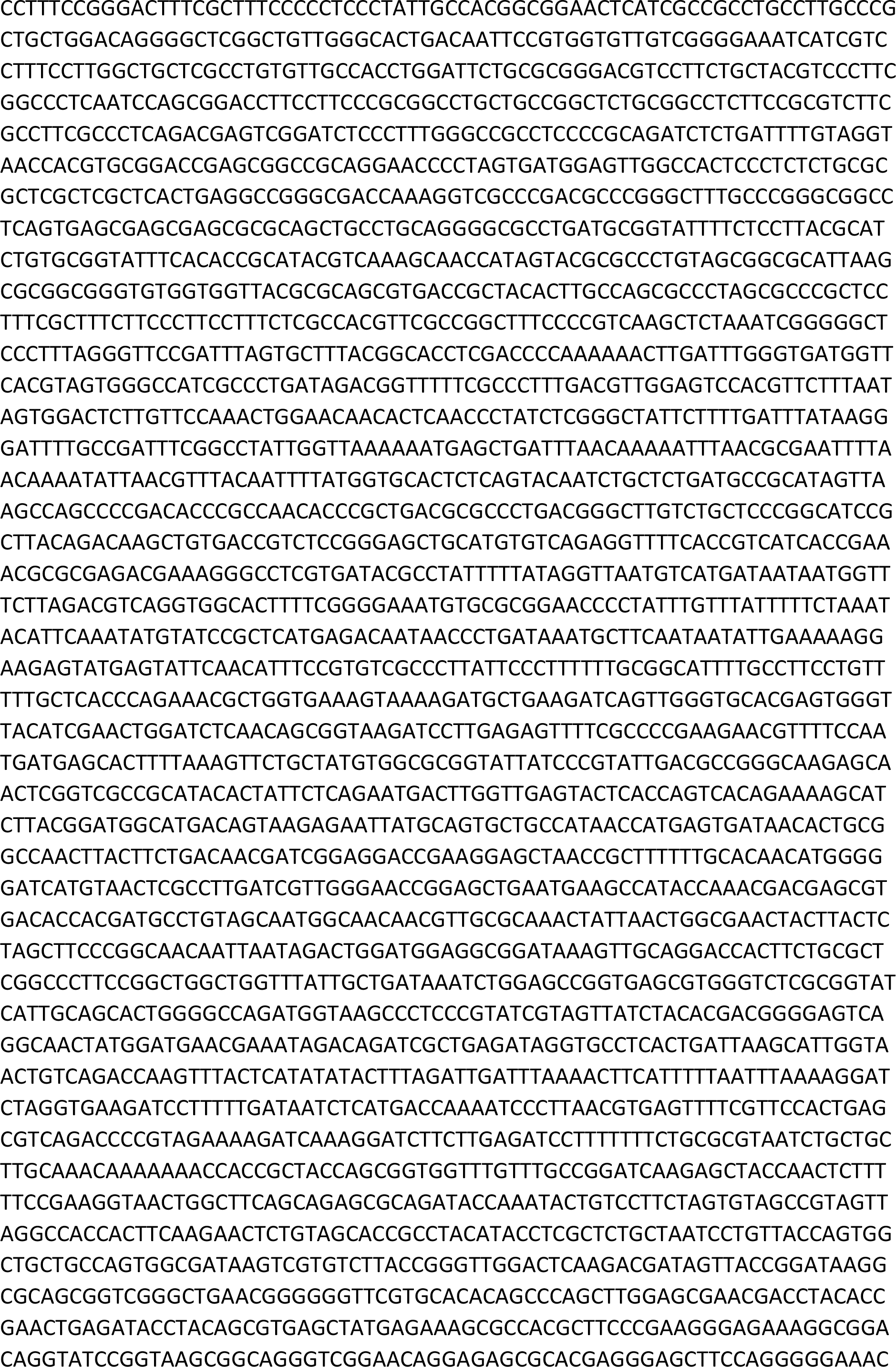
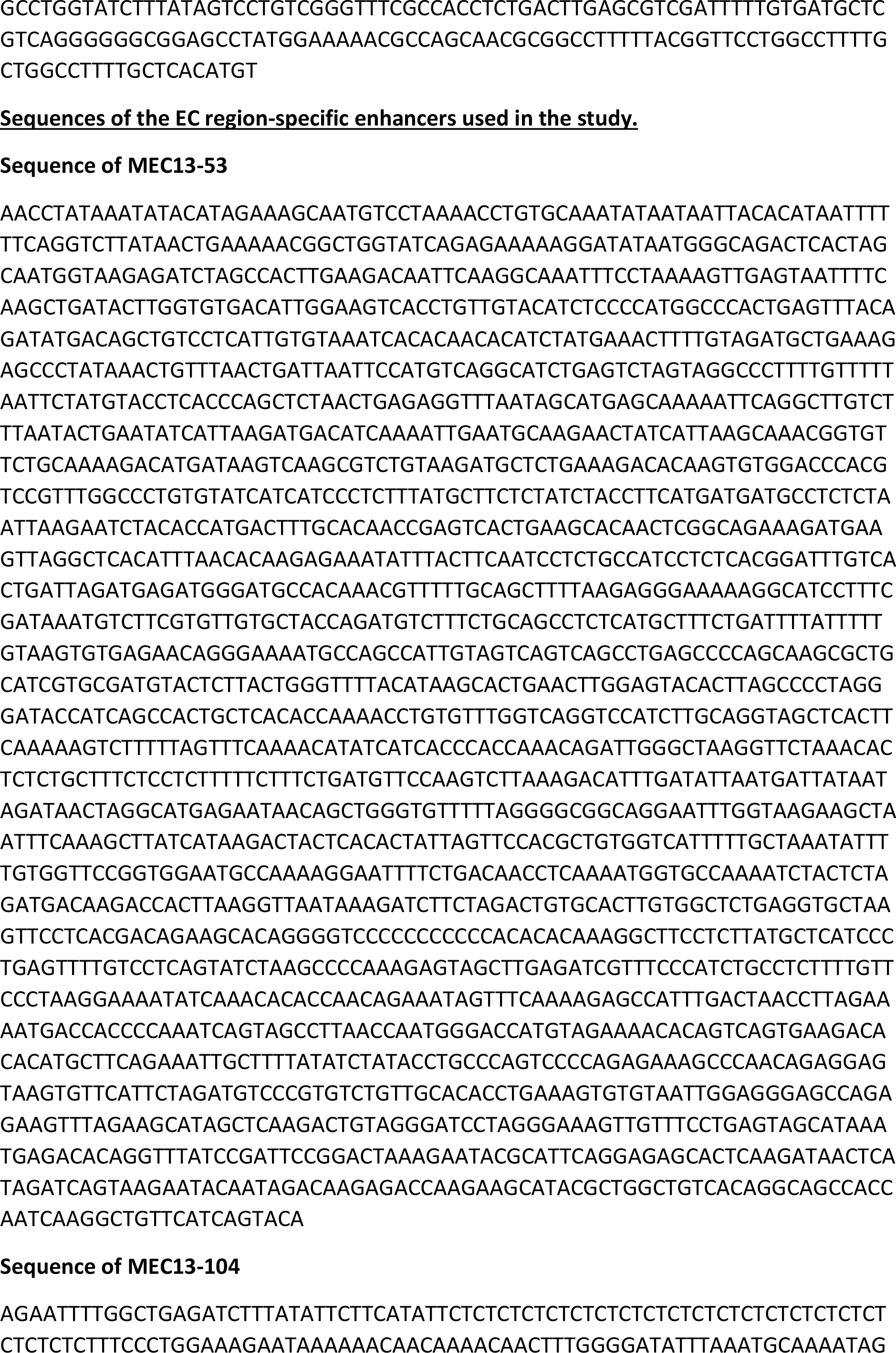
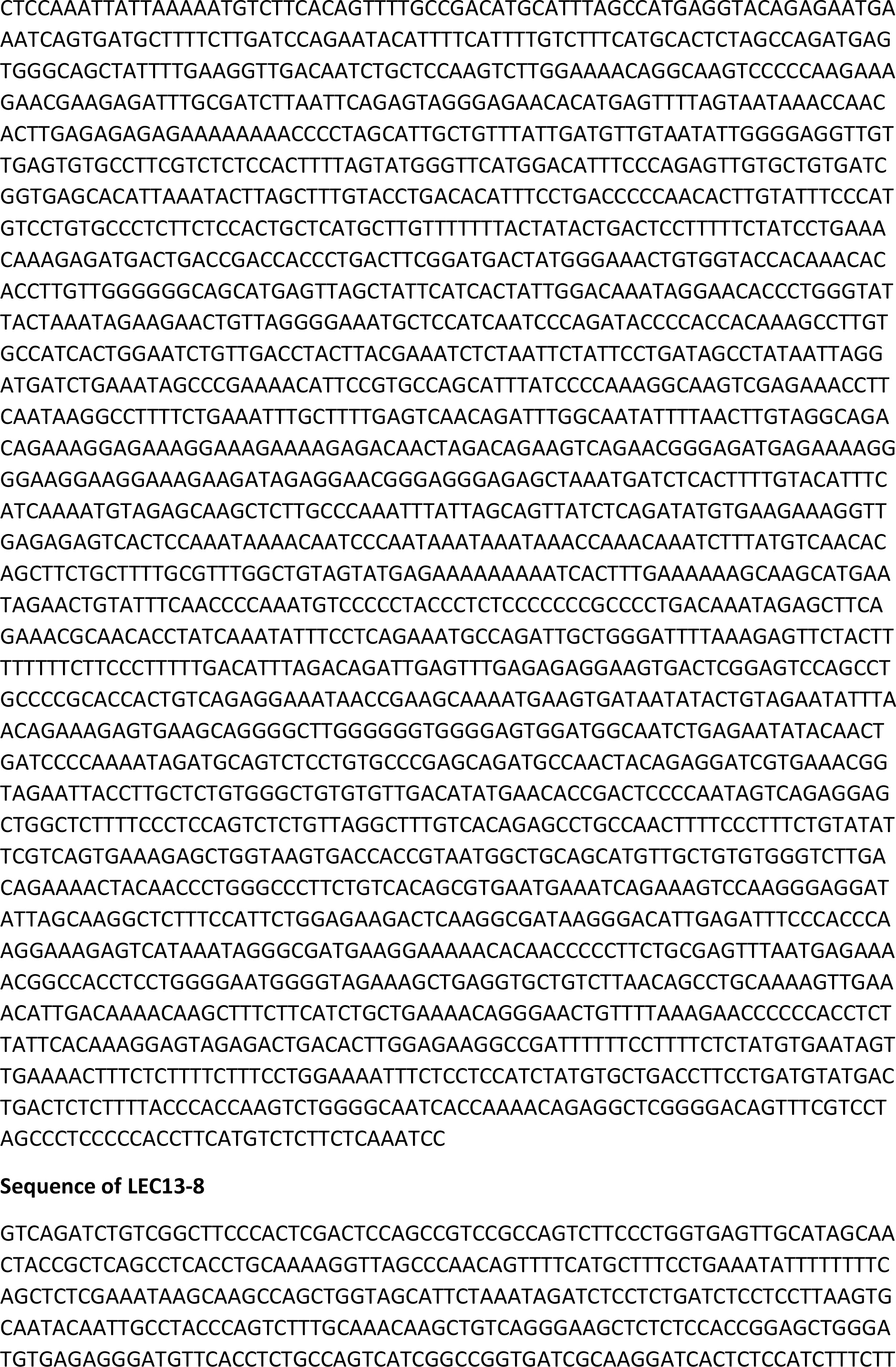
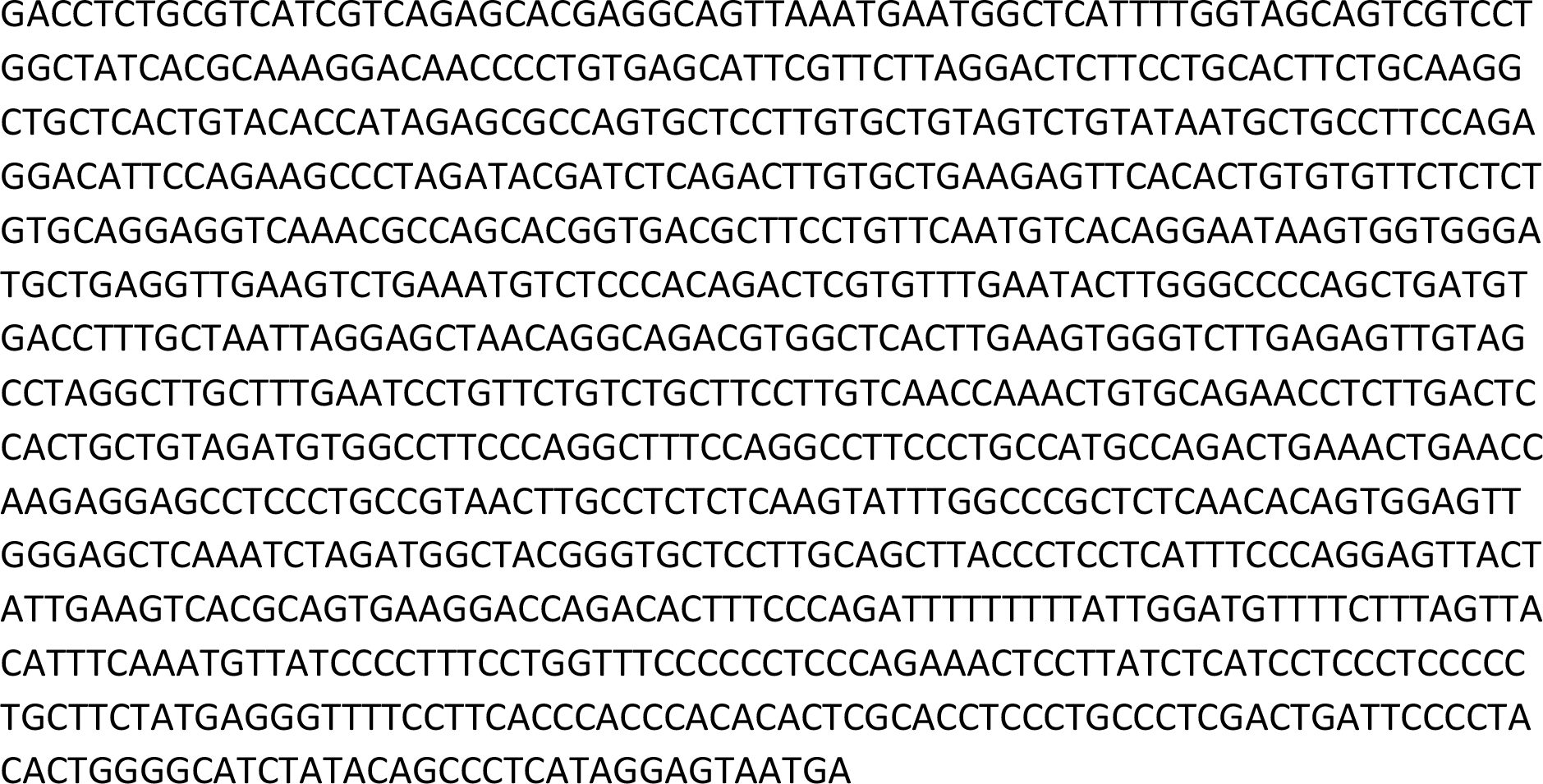

## Abbreviations

AD: Alzheimer’s disease
CB: Calbindin
CMV: Cytomegalovirus
Dlx: *Distalless*
EC: Entorhinal cortex
EDGE: Enhancer Driven Gene Expression
FGF4: Fibroblast growth factor 4
GFP: Green fluorescent protein
ITR: Inverted terminal repeats
LEC: Lateral entorhinal cortex
LII: Layer II
MEC: Medial entorhinal cortex
rAAV: Recombinant Adeno-associated virus
RE: Reelin
TK: Thymidine kinase
TVAG: Tumor virus receptor A (Avian) and Rabies glycoprotein

## Acknowledgement

The work was funded by FRIPRO ToppForsk grant Enhanced Transgenics (90096000) of the Research Council of Norway, the Kavli Foundation, the Centre of Excellence scheme of the Research Council of Norway-Centre for Biology of Memory and Centre for Neural Computation, The Egil and Pauline Braathen and Fred Kavli Centre for Cortical Microcircuits, and the National Infrastructure scheme of the Research Council of Norway-NORBRAIN.

We acknowledge the help from Christina Schrick and Qiangwei Zhang for immunostainings and In situ hybridizations, Grethe M. Olsen and Miguel Carvalho for training on rat injections, Nicola P. Montaldo for demonstrating qPCR, Hanne M. Møllergård, Siv Eggen and technicians at the animal facilty (Kavli institute/CNC), Ben Kanter and Christine M. Lykken for valuable suggestions on the manuscript, Tomas Bjorklund and Menno P. Witter and members of Kentros lab for helpful discussions.

