## Supplementary figures and images for "Generation of viral vectors specific to neuronal subtypes of targeted brain regions by Enhancer-Driven Gene Expression (EDGE)"

### Supplemental figures

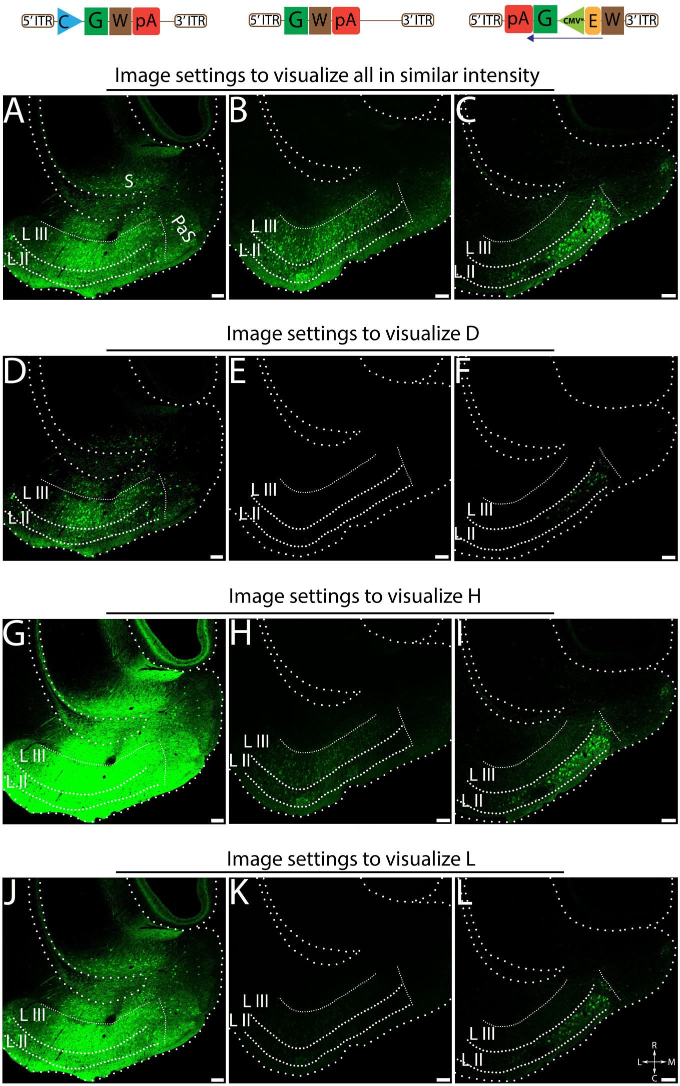

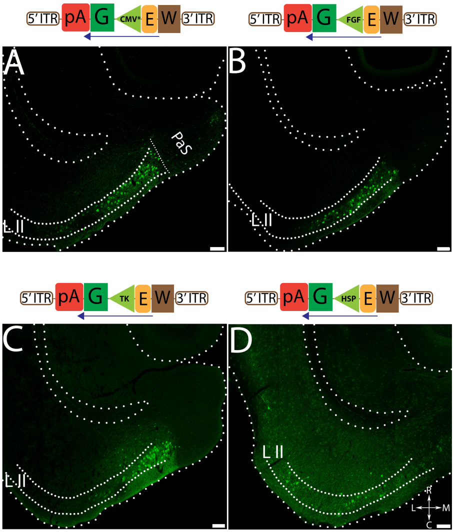

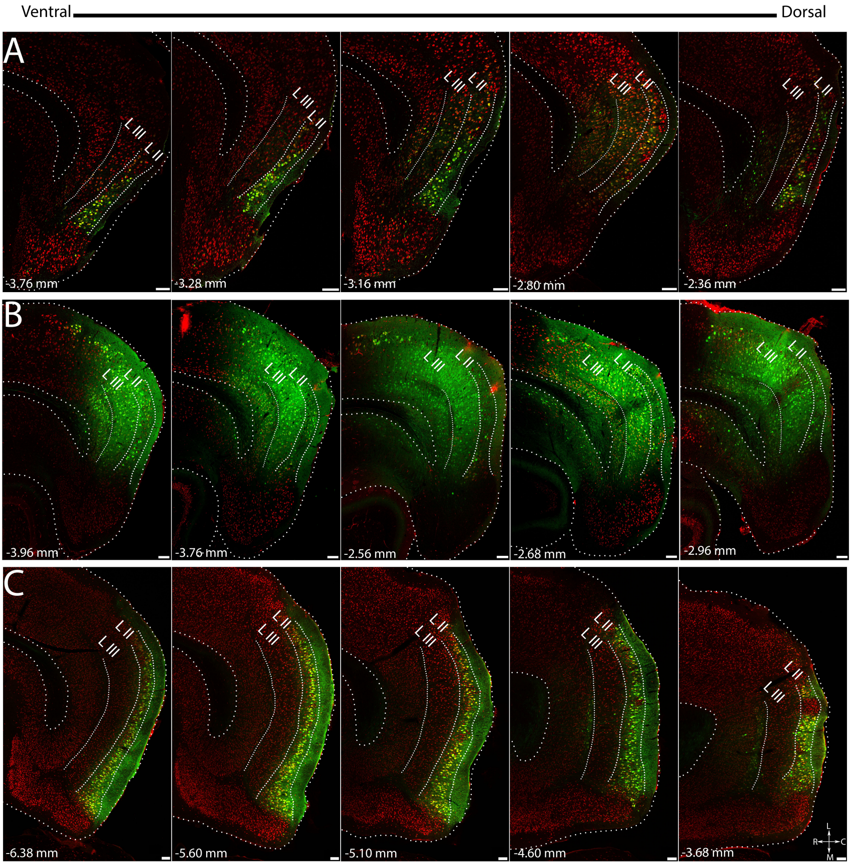
